# Mechanism of Mg^2+^ Mediated Encapsulation of an Anionic Cognate Ligand in a Bacterial Riboswitch

**DOI:** 10.1101/2022.05.19.492696

**Authors:** Sunil Kumar, Govardhan Reddy

## Abstract

Riboswitches in bacteria regulate gene expression and are targets for antibiotic development. The fluoride riboswitch is essential for bacteria’s survival as it is critical to maintaining the F^−^ ion concentration below the toxic level. The anionic cognate ligand, F^−^ ion, is encapsulated by three Mg^2+^ ions in a trigonal pyramidal arrangement bound to the ligand-binding domain (LBD) of the riboswitch. The assembly mechanism of this intriguing LBD structure and its role in transcription initiation are not clear. Computer simulations using both coarse-grained and all-atom RNA models show that F^−^ and Mg^2+^ binding to the LBD are essential to stabilize the LBD structure and tertiary stacking interactions. We propose that the first two Mg^2+^ ions sequentially bind to the LBD through water-mediated outer-shell coordination. The first bound Mg^2+^ should undergo a transition to a direct inner shell interaction through dehydration to strengthen its interaction with LBD before the binding of the second Mg^2+^ ion. The binding of the third Mg^2+^ and F^−^ to the LBD occurs in two modes. In the first mode, the third Mg^2+^ binds first to the LBD, followed by F^−^ binding. In the second mode, Mg^2+^ and F^−^ form a water-mediated ion pair and bind to the LBD simultaneously, which we propose to be the efficient binding mode. We show that the linchpin hydrogen bonds involved in the antiterminator helix formation and transcription initiation are stable only after F^−^ binding. The intermediates populated during riboswitch folding and cognate-ligand binding are potential targets for discovering new antibiotics.

## Introduction

Riboswitches are noncoding RNAs that modulate gene expression in response to cognate ligand binding. Cations like Mg^2+^, Na^+^, and K^+^ condense on the riboswitch to counter-balance the negative backbone charge and stabilize the folded conformations relevant for its function. An anionic ligand has to breach the positively charged cloud of cations to approach and interact with the RNA. Further, the polyanionic nature of RNA excludes anions from its ionic atmosphere,^1,2^ which makes the study of anionic ligand binding to RNA interesting and challenging.

Fluoride sensing riboswitch is an exciting example of RNA-anion interaction where the smallest biologically relevant anion, F^−^, acts as a cognate ligand and specifically binds to RNA through the cationic pocket composed of three Mg^2+^ ions. The fluoride riboswitch aptamer domain (FAD) has unparalleled selectivity towards only F^−^ among all the halide anions.^3^ Further, the presence of fluoride riboswitch in 11 human bacterial pathogens^4^ makes it an important target in the development of new antimicrobials^5,6^ as this riboswitch is critical for maintaining the cytoplasmic F^−^ concentration ([F^−^]) in the bacteria below the toxic levels.

We studied the mechanism of folding and fluoride ion encapsulation by the three Mg^2+^ bound to the ligand binding domain (LBD) of fluoride riboswitch aptamer domain (FAD) from *crcB* gene of *Thermotoga petrophila*. The FAD is 52 nt long, and in the native state (*holo*-form), it has the following tertiary structural elements (TSE) helical stems S_1_ and S_2_ joined by an internal loop that interacts with the 5′ nucleotides to form a pseudoknot (PK)^3^ (Figure 1). Three Mg^2+^ bind to the LBD and create a cationic pocket where F^−^ binds with high specificity. LBD comprises outward-pointing phosphate groups of five nucleotides: A6, U7, G8, U41 and G42 (LBD-nucleotides). FAD has six tertiary stacking interactions in the vicinity of the LBD to hold the backbone of LBD-nucleotides in a position conducive for Mg^2+^ and F^−^ binding and enhance its stability (Figure S2).

**Figure 1:**
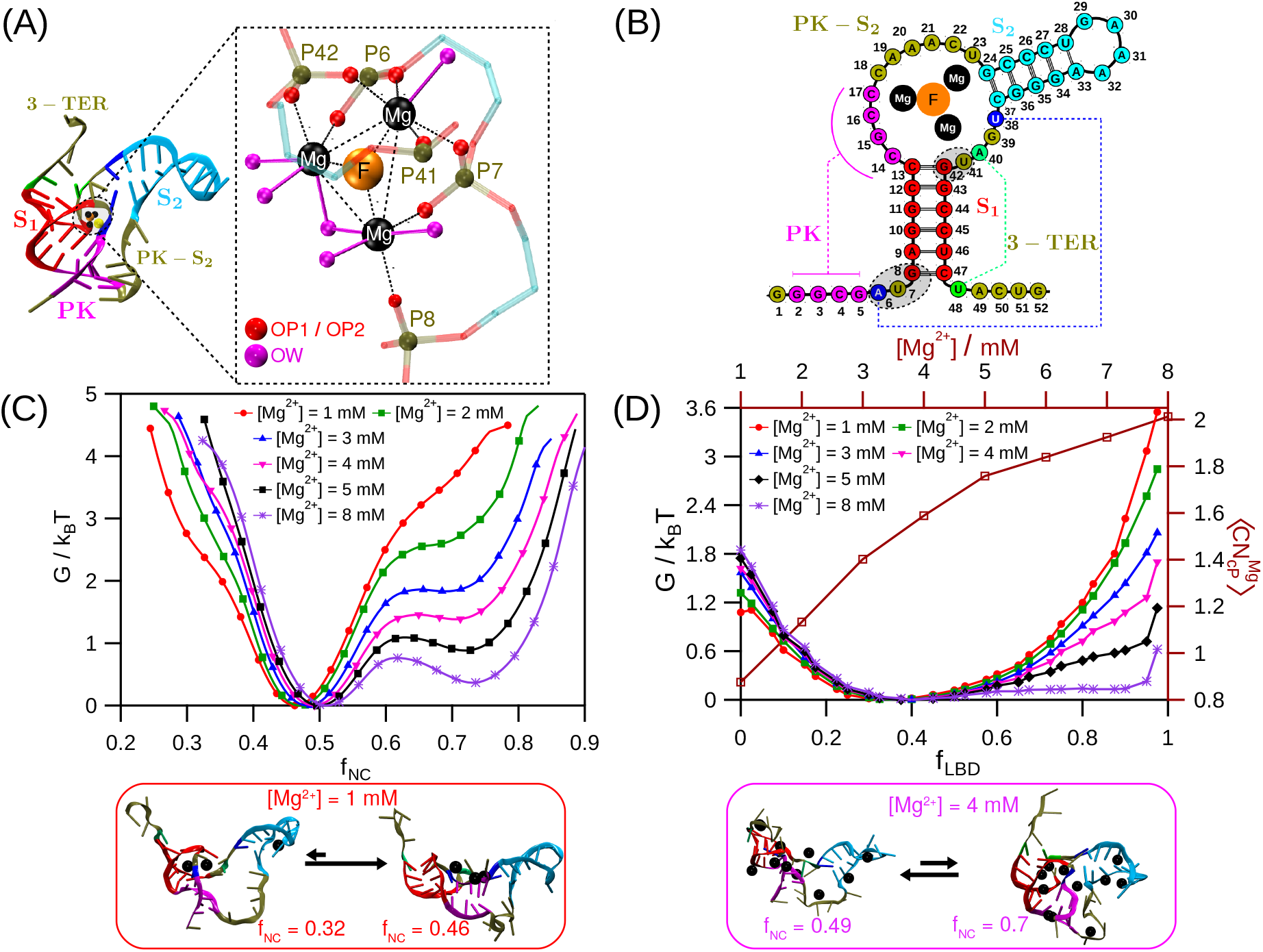
(A) Crystal structure of the fluoride aptamer domain (FAD) (PDB: 4ENC) ^3^ shown in cartoon representation. The tertiary structural elements - pseudoknot (PK), helical stem-1 (S_1_) and stem-2 (S_2_) are shown in magenta, red and cyan, respectively. The linchpin hydrogen bond pairs A6/U38 and A40/U48 are shown in blue and green cartoon representation, respectively. The nucleotides located between PK and S_2_ (referred as PK-S_2_), and at the 3′ terminal (referred as 3-TER) are shown in tan. Mg^2+^ and F^−^ are shown as black and orange beads, respectively. The K^+^ near Mg^2+^ is shown as yellow bead. The three Mg^2+^ interact with the O atoms of the phosphate group of nucleotides A6, U7, G8, U41 and G42 shown in gray shaded area in the schematic. The ligand binding domain (LBD) is highlighted inside the dotted rectangle. Backbone of the five nucleotides A6, U7, G8, U41 and G42 (referred as LBD-nucleotides) are shown with translucent stick model. The LBD forming RNA atoms (referred as LBD RNA-atoms) include two phosphate group oxygen atoms (OP) each from A6, U7 and G42, and one OP atom each from G8 and U41. All the LBD RNA-atoms are shown as red beads. All the octahedral coordinating sites for each of the Mg^2+^ are occupied either by the phosphate O atom (shown as red beads) or water O atom (shown as magenta beads). The P atoms of the LBD-nucleotides are shown as tan beads. The three Mg^2+^ which are bound to LBD (referred as LBD-3Mg) are located at the octahedral holes created by the LBD RNA-atoms and water molecules. The LBD-3Mg constitute the trigonal base of the cationic pocket and F^−^ occupies the fourth vertex of the trigonal pyramidal unit. (B) Two dimensional structure of the FAD is shown using circle and stick representation. The nucleotides regions are marked using the similar coloring scheme as mentioned before. The linchpin hydrogen bond pairs A6/U38 and A40/U48 are shown in blue and green circles, respectively.(C) Free energy surface (FES) of FAD is projected onto fraction of native contacts, *f*_NC_. FAD populates two major basins corresponding to the dominant unfolded and folded states. A sparsely populated unfolded state is observed in low [Mg^2+^]. Representative conformation of the FAD in each basin of the FES is shown in cartoon representation. The equilibrium between conformations in different basins are shown for [Mg^2+^] = 1 and 4 mM. (D) FES of the FAD (left axis) is projected onto the fraction of LBD native contacts, *f*_LBD_ (bottom axis), for different [Mg^2+^]. The coordination number (right axis) of the Mg^2+^ with respect to the LBD-nucleotide phosphate beads, 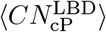 as a function of [Mg^2+^] (top axis).

Alkali and alkaline earth metal ions are essential to the folding of RNA. Experiments^7–20^ and computer simulations^21–26^ probed the effect of various cations and cognate ligands on the folding kinetics and thermodynamics of multiple riboswitches. The ion-counting experiment demonstrated the preferential accumulation of cations and exclusion of anions from the ionic atmosphere of RNA.^1^ Single molecule FRET study on P4-P6 domain of the *Tetrahymena* group I intron carried out to probe the RNA-anion interactions, demonstrated that anion identity can affect the RNA folding thermodynamics and kinetics.^27^ SHAPE-seq,^28^ CEST-NMR,^29^ and smFRET^30^ studies of the *Bacillus cereus crcB* fluoride riboswitch showed that in the absence of F^−^ binding, the aptamer can populate both the *holo*-like structure and the unfolded state. The *holo*-like structure stabilizes only upon F^−^ binding to the FAD suggesting that fluoride riboswitch follows conformational selection mechanism for ligand binding. SHAPE-seq^28^ and CEST-NMR^29^ studies also suggested that the nucleotides located in between the pseudoknot (PK) and the helix S_2_, and the 3′ tail can be a potential indicator to differentiate between the unfolded and folded state of the aptamer (Figure 1A,B). CEST-NMR study^29^ on the FAD of *Bacillus cereus* suggested that in the presence of Mg^2+^, there exists a sparsely populated *apo*-excited state with *holo*-like conformation but with ruptured “linchpin” hydrogen bond in the absence of F^−^ binding. The ruptured linchpin hydrogen bond enables the formation of terminator helix which in turn signals the transcription termination. Single-molecule FRET^30^ study demonstrated that the hydrogen bond between nucleotides A40/U48 forms and ruptures upon F^−^ ion binding and unbinding, respectively. Computations using both static and dynamic DFT calculations^31^ probed the structural stability of the cluster formed by the three Mg^2+^, one fluoride, phosphates of LBD, and water in the FAD of *Thermotoga petrophila*. DFT results show that the cluster is stable in itself, and once assembled, it does not require any additional stabilization from the overall RNA.

It is highly challenging for any experimental technique to simultaneously track the folding of various secondary and tertiary structural elements in RNA and probe the contribution of multiple ions to the stability of these structures. Molecular dynamics (MD) simulations have the potential to study ion assembly and RNA folding simultaneously. However, it is not trivial even for MD simulations to probe the folding and ion assembly in systems such as fluoride riboswitch as multiple time scales ranging from nanoseconds to seconds are involved. To overcome this problem in MD simulations, we studied the FAD folding and F^−^ binding using both coarse-grained and all-atom models of RNA. Using coarse-grained models, we show that in the presence of Mg^2+^, FAD attains *holo*-form like structure with similar tertiary structural elements irrespective of its cognate ligand, F^−^ binding. However, F^−^ binding is imperative for the stabilization of the LBD and the linchpin hydrogen bond. Using all-atom simulations, we probed the mechanism of F^−^ encapsulation by the three Mg^2+^ bound to the LBD. We provide evidence that the three Mg^2+^ bind to the LBD in a sequential manner. For efficient binding, the third Mg^2+^ and the F^−^ approach the LBD as a water-mediated ion pair. The all-atom simulations further show that the linchpin hydrogen bond between A40 and U48, which plays a role in the formation of terminator helix preventing transcription, is unstable in the absence of F^−^ binding.

## Methods

We studied the effect of Mg^2+^ on the folding of Fluoride riboswitch aptamer domain (FAD) from *crcB* gene of *Thermotoga petrophila* using the coarse-grained three interaction site (TIS) RNA model^32,33^ and Langevin dynamics simulations. We performed all-atom molecular dynamics simulations to understand the mechanism of Mg^2+^ mediated F^−^ sensing by the FAD.

### Simulations

#### TIS Model of RNA

The TIS model for FAD is constructed using the crystal structure (PDB: 4ENC).^3^ In the TIS model^32,33^ each nucleotide is represented using three sites representing the phosphate, sugar, and base groups. The positions of phosphate, sugar, and base sites are the centers of mass of their respective chemical groups. The monovalent (K^+^ and F^−^) and divalent (Mg^2+^) ions are explicitly present in the simulation. The Hamiltonian for the TIS model is given by

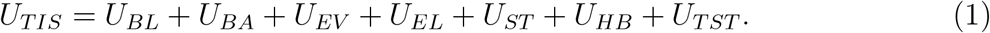

The different components in the Hamiltonian correspond to the bond length (*U*_BL_), bond angle (*U*_BA_), excluded volume repulsion between different sites (*U*_EV_), electrostatic interaction between charged sites (*U*_EL_), single-strand base stacking interaction among consecutive bases (*U*_ST_), hydrogen bonding interaction (*U*_HB_) and tertiary stacking interaction between two non-consecutive bases (*U*_TST_), respectively. The force field details and parameters for RNA sites can be found in the works of Denesyuk and Thirumalai.^32,33^

The list of base pairs involved in the native hydrogen bond network present in the folded structure of FAD is obtained using the crystal structure (PDB: 4ENC) and WHAT IF web server (https://swift.cmbi.umcn.nl). ^34^ The base pairs with a hydrogen bond between them interact using the hydrogen bonding potential (*U*_HB_). The model takes into account the tertiary base stacking interactions present in the crystal structure. The TIS-RNA model allows the formation of non-native canonical hydrogen bonds but not the formation of non-native non-canonical hydrogen bonds. ^33^ In this study we have not allowed the formation of any non-native hydrogen bonds. This model is successful in quantitatively accounting for the folding thermodynamics of multiple RNA systems^26,33,35,36^ demonstrating that it is reliable and transferable to study various other RNA systems.

#### Coarse-Grained Simulations

Coarse-grained simulations are performed to study the role of Mg^2+^ in the folding of FAD as [Mg^2+^] is varied from 1 mM to 8 mM. The [K^+^] is fixed at 30 mM in all the simulations. The simulations are performed in a cubic box of length 165 Å. The number of Mg^2+^ and K^+^ ions in the simulation box are computed using their concentration and box volume. Only the cognate ligand for the FAD, the F^−^ ions, are added to maintain charge neutrality in the simulation box. [F^−^] increases from 13 mM to 27 mM with the increase in [Mg^2+^] from 1 mM to 8 mM. All ions are explicitly modeled as beads with charge and excluded volume. The excluded volume force field parameters for K^+^, F^−^, and Mg^2+^ ions are taken from the modified AMBER ff14 force field. ^37^ We used Langevin dynamics simulations to study the folding dynamics of FAD at temperature, *T* = 310 K. To compute the thermodynamic properties of FAD folding, we used 5% viscosity of water, *η* = 5 *×* 10^−5^ Pa·s to enhance the conformational sampling rate. The equation of motion for the RNA sites and ions in Langevin dynamics is given by

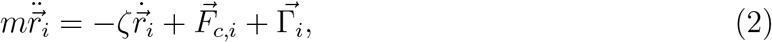

where *m*_*i*_ is the mass (in amu), *ζ*_*i*_ (= 6*πηR*_*i*_) (in amu/fs) is the friction coefficient, 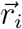 is the position, 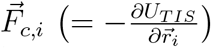 is the deterministic force, *R*_*i*_ is the radius (in Å) of *i*^*th*^ site in the system. 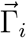 is the random force on the *i*^*th*^ site with a white-noise spectrum. The random force auto-correlation function is given by 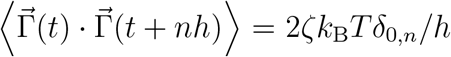, where *n* = 0, 1, …, *δ*_0,*n*_ is Kronecker delta function, and *k*_B_ is the Boltzmann constant. The Langevin equation is integrated using the velocity Verlet algorithm with a time step *h* (= 2.5 fs).^33,38^ System coordinates are saved after every 5,000 steps (*τ*_*f*_ = 12.5 ps) to compute the properties. The initial 0.5 *μ*s of simulation data is ignored in computing the properties. For each [Mg^2+^], at least ≈ 7 *μ*s of simulation data is collected to compute the average properties. We used VMD to generate the three dimensional structures of the FAD.^39^ The atomistic coordinates of the FAD are generated using the coarse-grained coordinates and TIS2AA^40^ program, which uses the fragment-assembly approach^41^ and energy minimization embedded in AmberTools.^42^

#### All-Atom Simulations

We started the all-atom simulations of FAD using the crystal structure. We performed all-atom simulations for five different FAD systems in the presence and absence of crystal bound Mg^2+^ and F^−^ (Table 1). MD simulations were performed using GROMACS-5.1.8^43,44^ with modified AMBER ff14 force field^37^ for RNA and TIP4P-D water model.^45^ Initially, FAD was placed at center of a cubical box of length 85 Å, and solvated with TIP4P-D water molecules. The required number of Mg^2+^, K^+^, F^−^ and Cl^−^ ions were added to the simulation box (Table 1). The energy of the system is minimized using steepest descent algorithm with a maximum force tolerance of 1000.0 kJ mol^−1^nm^−1^. The energy minimized system was equilibrated in the following steps: (1) The positions of RNA and ions were kept fixed using harmonic position restraints with a spring constant of 1000.0 kJ mol^−1^nm^−2^ and water molecules were allowed to equilibrate in the NVT ensemble for 10 ns. This step ensured that hydration shells are formed for all the ions. (2) Position restraints were applied only on RNA atoms, whereas ions and water molecules are allowed to equilibrate in the NVT ensemble for 10 ns so that the hydrated ions condense around the negatively charged RNA. (3) Finally, position harmonic restraints were eliminated on all the atoms and the system is allowed to equilibrate in the NPT ensemble for 10 ns.

**Table 1:** Details of various all-atom simulation systems.

| System | $N_{\text{Mg}}^{\text{cry}}$ | $N_{\text{F}}^{\text{cry}}$ | $N_{\text{Mg}}^{\text{bulk}}$ | $N_{\text{F}}^{\text{bulk}}$ | $N_{\text{K}}$ | $N_{\text{Cl}}$ | [Mg<sup>2+</sup>]<br>(mM) | [F<sup>-</sup>]<br>(mM) | [K<sup>+</sup>]<br>(mM) | [Cl<sup>-</sup>]<br>(mM) | Cumulative<br>Simulation Time ($\mu\text{s}$) |
| --- | --- | --- | --- | --- | --- | --- | --- | --- | --- | --- | --- |
| AA[$\phi$] (*apo*) | 0 | 0 | 0 | 0 | 51 | 0 | 0 | 0 | 139 | 0 | $\approx 4$ |
| AA[0,0] | 0 | 0 | 20 | 20 | 55 | 24 | 55 | 55 | 150 | 66 | $\approx 15$ |
| AA[1,0] | 1 | 0 | 19 | 20 | 55 | 24 | 55 | 55 | 150 | 66 | $\approx 16$ |
| AA[2,0] | 2 | 0 | 18 | 20 | 55 | 24 | 55 | 55 | 150 | 66 | $\approx 14$ |
| AA[3,1] (*holo*) | 3 | 1 | 17 | 19 | 55 | 24 | 55 | 55 | 150 | 66 | $\approx 5$ |

$N_{\text{Mg}}^{\text{cry}}$ and $N_{\text{F}}^{\text{cry}}$ are the number of bound Mg<sup>2+</sup> and F<sup>-</sup> to the LBD. $N_{\text{Mg}}^{\text{bulk}}$ and $N_{\text{F}}^{\text{bulk}}$ are the number of Mg<sup>2+</sup> and F<sup>-</sup> in the simulation box and away from the LBD. $N_{\text{K}}$ and $N_{\text{Cl}}$ are the number of K<sup>+</sup> and Cl<sup>-</sup> in the simulation box. We ran four independent trajectories for system with AA[0,0] configuration. We ran two independent simulations for the rest of the system configurations.

The temperature *T* of the system is maintained at 310 K using a modified Berendsen thermostat^46^ with a relaxation time *τ*_T_ = 0.1 ps. The pressure of the system is maintained at 1 atm using Parrinello-Rahman barostat^47^ with *τ*_*P*_ = 2.0 ps and isothermal compressibility of water. We have used 10 Å as cut-off for both Lennard-Jones and Coulombic interactions. Long-range Coulombic interactions were computed using Particle Mesh Ewald (PME)^48^ sum method with a grid space of 1.6 Å. Long-range dispersion correction for energy and pressure were also incorporated. All bonds involving hydrogen atom were constrained using LINCS algorithm^49^ throughout the production stage of the simulation. For all the five systems, we performed two independent simulations and the cumulative simulation time is reported in Table 1.

## Data Analysis

### Fraction of Helix Formation

The average fraction of helix formation for a given helix *H*, is computed using the equation

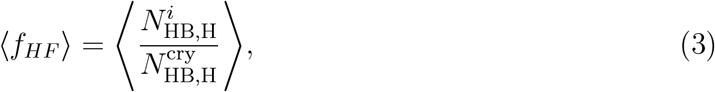

where 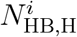 and 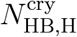 are the number of hydrogen bonds present in the helix *H* in *i*^*th*^ conformation and crystal structure, respectively. ⟨ ⟩ denotes the average over all the conformations. A hydrogen bond is considered to be present if its energy is lower than the thermal energy (*k*_B_*T*).^33^ Similarly, the linchpin hydrogen bonds A6/G38 and A40/U48 defined to be formed if the hydrogen bond interaction energy is less than *k*_B_*T*.

### Local [Mg^2+^] Around Phosphate Sites

The local [Mg^2+^] in the vicinity of the phosphate site of the *i*^*th*^ nucleotide in molar units is computed^50^ using the relation

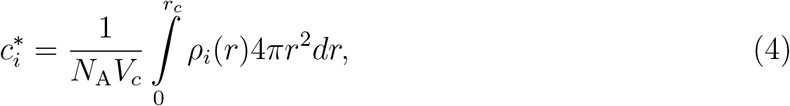

where *ρ*_*i*_(*r*) is the number density of the Mg^2+^ at a distance *r* from the *i*^*th*^ phosphate site, *V*_c_ is the spherical volume of radius *r*_*c*_, and *N*_A_ is the Avogadro’s number. The cutoff radius *r*_c_ is given by *r*_c_ = *R*_Mg_ + *R*_P_ + Δ*r* ≈ 5 Å where *R*_Mg_ (1.185 Å) and *R*_P_ (2.1 Å) are the radii of Mg^2+^ and phosphate sites, and Δ*r* is the margin distance. We used Δ*r* = 1.7 Å to ensure that we take into account only the tightly bound local Mg^2+^ around the phosphate sites. We computed 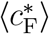 using the similar equation as Eq. 4 with *r*_c_ = *R*_P_ + *R*_Mg_ + *R*_F_ + Δ*r* ≈ 7.3 Å, where *R*_F_ (2.3 Å) is the radius of F^−^ site.

### Free Energy Surface (FES) Calculation

The total number of native contacts in the FAD is computed using the TIS model of the folded crystal structure (PDB: 4ENC).^3^ A pair of sites *i* and *j* in the crystal structure are defined to have a native contact between them if |*i* − *j*| > 10, and the distance between the sites *r*_ij_ is less than 15 Å (see SI Figure S1). The *f*_NC_ for *i*^*th*^ FAD conformation is computed using the equation

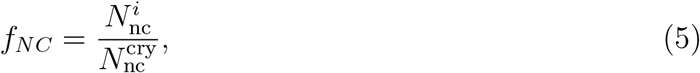

where 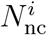 is the number of native contacts present in the *i*^*th*^ conformation and 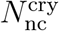 is the number of native contacts present in the crystal structure.

The FES corresponding to the FAD folding (*G*) is projected onto the fraction of native contacts *f*_NC_, and it is calculated using the equation

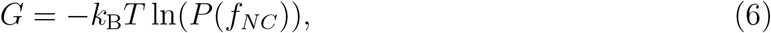

where *P*(*f*_NC_) is the probability distribution of *f*_NC_.

To probe the role of C18 - U23 nucleotides located between PK and S_2_ (PK-S_2_ nucleotides) in FAD folding we computed 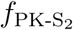, fraction of native contacts between the PK-S_2_ nucleotides and their neighboring nucleotides (labelled as 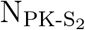) that are located within 15 Å from PK-S_2_ and at least three nucleotides away along the RNA contour in the FAD crystal structure. Similarly, to probe the role of A49 - G52 nucleotides located at the 3′-terminal (3-TER nucleotides) we computed *f*_3-TER_, fraction of native contacts between the 3-TER nucleotides and their neighboring nucleotides (labelled as N_3-TER_) that are located within 15 Å from 3-TER and at least three nucleotides away along the RNA contour in the FAD crystal structure.

### Coordination Number Calculation

We computed the coordination number 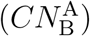 of particle *i* belonging to group A with the particle *j* belonging to group B in a FAD conformation using the COORDINATION module of PLUMED library (v2.6.4).^51,52^ The 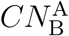 is computed for every conformation using the following relation,

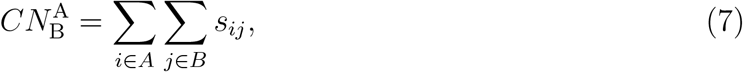

where *s*_*ij*_ is a switching function and is given by

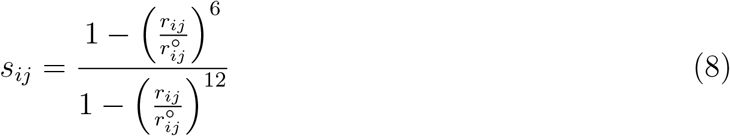

where *r*_*ij*_ is the distance between *i*^*th*^ particle of group A and *j*^*th*^ particle of group B. 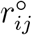 is the cut-off distance large enough to effectively capture the interaction between the particles *i* and *j*.

To account for the average number of Mg^2+^ bound to the LBD-nucleotides in the CG simulations for a given [Mg^2+^], we computed, 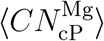, which is the average coordination number of Mg^2+^ with the center of mass of phosphate beads of LBD-nucleotides. We designated all the Mg^2+^ present in the simulation box for a given [Mg^2+^] as group A, and the center of mass of phosphate beads of the LBD-nucleotides (A6, U7, G8, U41, and G42) as group B. The cut-off distance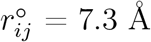, which is the Bjerrum length for water at 300 K.

To track the binding of F^−^ to the LBD in the all-atom simulations, we computed the coordination number individually for each of the F^−^ ion with the LBD RNA-atoms 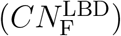. We designated the LBD RNA-atoms as group A, and a given F^−^ as group B. The cut-off distance, 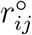 is 7.3 Å.

### Root Mean Square Deviation (RMSD) and Root Mean Square Fluctuation (RMSF)

We computed the RMSD of the conformations from the all-atom simulations with respect to the energy-minimized *holo*-form FAD crystal structure. We aligned all the conformations with respect to the energy-minimized *holo*-form structure of FAD before computing both RMSD and RMSF. To probe the changes in flexibility in the FAD structure due to the absence of ligand binding, we computed RMSF in the N1 atoms of all the FAD nucleotides, respectively (Figure S11B). Both RMSF and RMSD were computed using VMD.^39^ We used the simulation data from the systems AA[3,1] and AA[*ϕ*] for the analysis of *holo* and *apo* states of FAD, respectively.

### Mg^2+^ Binding Lifetime

To probe the mechanism of Mg^2+^ binding to the LBD of FAD, we computed the average conditional binding lifetime (*τ*(*i, j*)) and the occupancy time 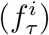 of the Mg^2+^ to the LBD in all-atom simulations. The Mg^2+^ within the binding cut-off distance (*R*_b_) from the phosphorus atoms (P atom) of all the LBD-nucleotides is labeled as bound to the LBD (Figure S9). The *R*_b_ is 11 Å for the AA[0,0] and AA[1,0] and 10 Å for AA[2,0]. The *R*_b_ value is estimated from the mean of average distances between P atoms of nucleotides A6/U41 and G8/G42.

### Dynamical Cross-Correlation Map (DCCM)

To probe dynamical cross-correlation^53^ between the nucleobases, we first coarse-grained the all-atom FAD conformations using the TIS-RNA bead description.^32,33^ Each nucleotide is described by phosphate (P), sugar (S) and nucleobase (B) beads and the position of these beads is at the center of mass of their respective chemical groups. The quantitative estimate for the correlated motion between a pair of beads mimicking nucleobases *i* and *j* is given by

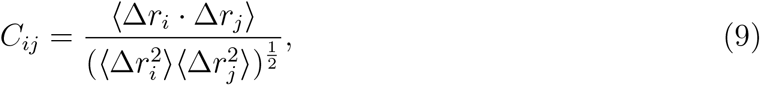

where Δ*r*_*i*_ is the displacement of *i*^*th*^ bead from its mean position. We used MD-TASK package^54^ to compute *C*_ij_. For a given pair of beads *i* and *j, C*_ij_ takes values between -1.0 to 1.0. The value -1.0 represents complete anti-correlation (beads move in opposite direction), the value 1.0 represents complete correlation (beads move in the same direction) and the value 0 represents no correlation (beads move in the perpendicular direction).

## Results

### LBD and Tertiary Stacking Interactions are Unstable in the Absence of F^−^ Binding

Noncoding RNA molecules are known to fold to a unique native state to perform gene regulation. Using the coarse-grained model of RNA^32,33^ and molecular dynamics simulations, we computed the free energy surface (FES) for the Mg^2+^ driven folding of FAD (see Eq. 6 in Methods) to probe the role of Mg^2+^ in facilitating F^−^ binding and corresponding conformational changes in FAD (Figure 1C). In computing the FES, the cognate ligand F^−^ is used as a counter ion to the Mg^2+^ to maintain charge neutrality in the simulation box. The FES projected onto the fraction of native contacts (*f*_NC_) (Eq. 6) shows that the aptamer is dominantly a two state folder, as it populated two major basins, the unfolded state (*f*_NC_ ≈ 0.46 - 0.51) and folded state (*f*_NC_ ≈ 0.70 - 0.73). As we increased [Mg^2+^] from 1 mM to 8 mM, the minima of the major unfolded basin shifted from *f*_NC_ ≈ 0.46 to ≈ 0.51 indicating that there is a compaction in the aptamer size in the major unfolded ensemble (Figure 1C).

In low [Mg^2+^] (≤ 4 mM), the aptamer sampled a sparsely populated unfolded state (*f*_NC_ ≈ 0.32) and a dominantly populated unfolded state (*f*_NC_ ≈ 0.46 - 0.49) (Figure 1C). The shoulder in the FES corresponding to the sparsely populated unfolded state disappeared for [Mg^2+^] > 4 mM. With the increase in [Mg^2+^] (> 3 mM), the aptamer starts populating the folded state (*f*_NC_ ≈ 0.7 - 0.73) separated by a barrier from the dominant unfolded state (*f*_NC_ ≈ 0.46). The relative population of the folded state increased with the increase in [Mg^2+^] from 3 mM to 8 mM (Figure 1C). However, the dominant unfolded state (*f*_NC_ = 0.46 - 0.51) remained the global minimum, irrespective of [Mg^2+^] in the simulations using the coarse-grained model of RNA. CEST-NMR^29^ and the smFRET^30^ studies also demonstrated that in presence of Mg^2+^ FAD dynamically populates two states with formed helices and pseudoknot; one of the state lacks the essential tertiary contacts whereas the other is similar to the folded state.

The tertiary structural elements (TSE) of the aptamer, the pseudoknot (PK) and helices (S_1_ and S_2_), are stable irrespective of the [Mg^2+^] (Figure 1A,S3). This is in agreement with the experiments,^29^ which showed that the aptamer in both the *apo* and *holo* forms has the same hydrogen bond network and similar tertiary structure (Figure S3). The stability of the TSE at low [Mg^2+^] implies that monovalent ions ([K^+^] = 30 mM) are adequate to stabilize the native-like tertiary structure of the aptamer.

In the simulations using the coarse-grained RNA model, the dominant unfolded state of the FAD is more stable compared to the folded state even in high [Mg^2+^] (Figure 1C) because we did not observe the encapsulation of F^−^ by three Mg^2+^ in the LBD as observed in the crystal structure^3^(Figure 1A). Due to the absence of interactions between the LBD nucleotides, the bound F^−^, and three Mg^2+^; we observed that there are dynamic transitions between locally folded and unfolded states in the following regions of the aptamer: (a) the LBD domain of the aptamer (Figure 1D), (b) tertiary contacts formed by the nucleotides present between the pseudoknot PK and helix S_2_, labeled as PK-S_2_ nucleotides (Figure 1A,B, and S4B,C), (c) tertiary contacts formed by the nucleotides located at the 3′ terminal labeled as 3-TER nucleotides (Figure 1A,B and S4D), (d) the six native tertiary stacking interactions present between the nucleotides: A6/G24, A6/G39, U7/G39, G8/A40, A40/A49, and G5/G42 (Figure S2, S12A).

The nucleotides A6, U7, G8, U41 and G42 form the LBD and fold into an orientation that forms the scaffold with phosphate groups pointing towards the center of the cavity, which facilitates the binding of three Mg^2+^ and F^−^ (Figure 1A). FAD crystal structure shows that three Mg^2+^ (LBD-3Mg) form the base of trigonal pyramidal LBD, and facilitate the binding of F^−^ at the fourth vertex of the pyramid to form a 3Mg-1F cluster^3^ (Figure 1A). The free energy for the LBD formation projected onto the fraction of native contacts among the LBD-nucleotides, *f*_LBD_ (see Eq. S1 in SI) shows that the aptamer populated conformations where the LBD-nucleotides are in *holo*-like orientation, which supports the cationic pocket formation for [Mg^2+^] ≥ 4mM (Figure 1D). Although the population of conformations with the folded LBD increased with the increase in [Mg^2+^], the state with the partially folded LBD is the most stable state. We hypothesize that a stable LBD domain requires the binding of three Mg^2+^ and F^−^ (Figure 1A). However in the simulations with the coarse-grained RNA model we observed the binding of at most two Mg^2+^ and no F^−^ binding in the LBD (Figure 1D). The average coordination number of the LBD-nucleotide phosphate beads with Mg^2+^, 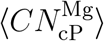 (see Methods), shows that the number of Mg^2+^ that bind to the LBD increased with [Mg^2+^], and approached the value 2 for [Mg^2+^] = 8 mM (Figure 1D).

These observations are in agreement with the NMR experiments^29^ on FAD, which have reported that the aptamer in *apo*-form without the bound F^−^ populates unfolded conformations where the pseudoknot PK, helices S_1_ and S_2_ are fully-formed but the local contacts around regions PK-S_2_ and 3-TER are ruptured (see SI for more details). These experiments further proposed that F^−^ binding is indispensable for the stability of the *holo*-like conformation. We did not observe a stable F^−^ binding in the coarse-grained simulations (see below). As a result, local unfolding is observed in multiple regions of the aptamer, and the unfolded state is the most stable state (Figure 1C).

### Mg^2+^ Ions Preferentially Bind to the LBD

To understand the role of Mg^2+^ in aptamer folding, we computed the average local Mg^2+^ concentration 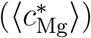 in the aptamer vicinity at nucleotide resolution (see Eq. 4 in Methods)^50^ using the data from coarse-grained simulations. Even at low [Mg^2+^], the ions are not randomly diffused along the RNA backbone. Mg^2+^ preferentially binds to the phosphate beads of LBD-nucleotides (A6 and G42) involved in the formation of RNA scaffold that facilitates the F^−^ binding (Figure 2A). For [Mg^2+^] ≤ 2 mM, Mg^2+^ only binds around the nucleotides G5 - G8 and U41 - G43. This observation is in agreement with the experiments,^3^ which demonstrated that Mg^2+^ binding in the vicinity of nucleotides A6 - G8 and U41 - G42 is essential for the cationic LBD formation. However, in the simulations using the coarse-grained RNA model, we observed at most two Mg^2+^ bind to the LBD nucleotides as indicated by the 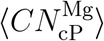 values (Figure 1D).

**Figure 2:**
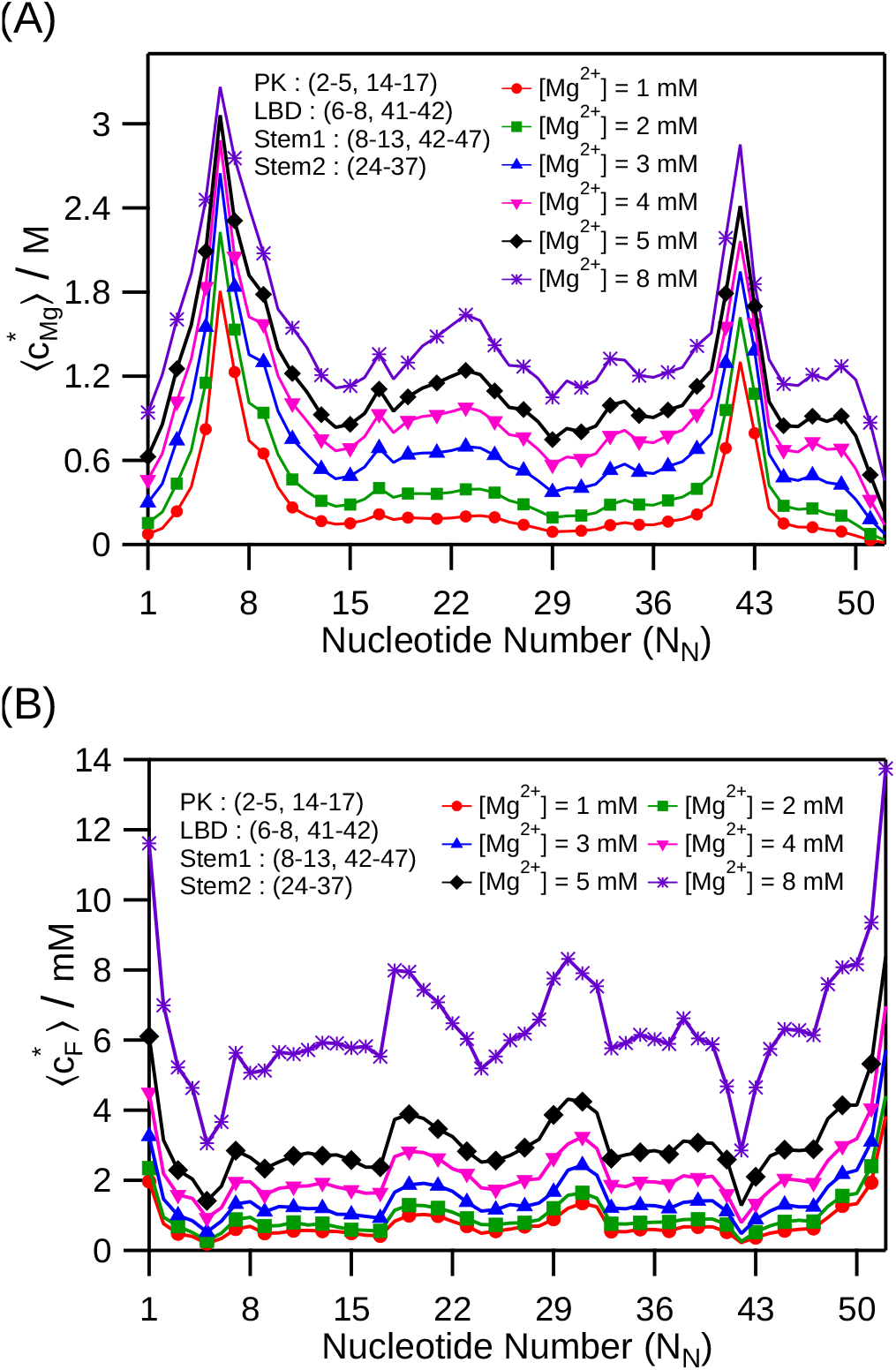
(A) The average local concentration of Mg^2+^ around individual P sites 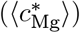 (in molar units) is plotted as a function of the nucleotide number (*N*_N_). The *N*_N_ range (starting from 1 as in the PDB ^3^) for the pseudoknot (PK), stems (S_1_ and S_2_) are in the annotation. For [Mg^2+^] = 1 mM, Mg^2+^ condenses around the LBD-nucleotides A6 and G42. As [Mg^2+^] increases, Mg^2+^ selectively accumulates around the same two nucleotides indicating that Mg^2+^ preferentially binds to these two nucleotides to create the cationic LBD. (B) The average local concentration of F^−^ around individual P sites 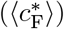 (in millimolar units) is plotted as a function of the nucleotide number (*N*_N_). The difference of three order in magnitude in the local concentrations of Mg^2+^ and F^−^ ions is due to excess cationic accumulation of Mg^2+^ and preferential exclusion of F^−^ from the vicinity of RNA, respectively.

In low [Mg^2+^] (≤ 2 mM), the lack of Mg^2+^ binding around the phosphate beads of PK-S_2_ nucleotides (*N*_N_ = C18 - U23) hinders the formation of tertiary contacts between PK-S_2_ and G1 - A6 nucleotides (part of the pseudoknot) due to the electrostatic repulsion (Figure S4C). Similarly, the lack of Mg^2+^ binding to the free 3-TER nucleotides (*N*_*N*_ = A49 - G52) prevents the formation of native contacts between 3-TER and U38 - U41 nucleotides, and also the formation of linchpin hydrogen bond A40/U48 (Figure S3 and S4A). With the increase in [Mg^2+^] (≥ 3 mM), Mg^2+^ binds in the vicinity of the nucleotides A19 - G24 (PK-S_2_), G5 - G8 and U41 - G43, and stabilizes FAD conformations with intact tertiary contacts between PK-S_2_ and G1 - A6 nucleotides, and local native contacts between 3-TER and U38 - U41 nucleotides (Figure S4C,D). However, even for [Mg^2+^] = 8 mM, the FAD conformations with the intact tertiary contacts are not the most stable state due to the absence of F^−^ binding to the LBD. The FAD state with fully-formed “linchpin” hydrogen bonding base-pairs, A6/U38 and A40/U48, which stabilize the native-like state and play a role in ligand binding signal transduction to the expression platform are not the most stable state (Figure S3).^3,29^ Although for [Mg^2+^] ≥ 5 mM, the FAD significantly populates folded state in absence of F^−^ binding event, implying [Mg^2+^] can drive the formation of conformations good enough to initiate the transcription. The CEST-NMR and single-round transcription assay studies^29^ also suggested that like the *holo*-form, the *apo*-form of the FAD with *holo*-like conformation can also activate transcription in presence of [Mg^2+^] ≥ 5 mM.

### F^−^ Transiently Binds to the LBD in the CG Simulations

The average local F^−^ concentration around the nucleotides 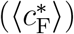 shows that unlike Mg^2+^, F^−^ are diffused around the RNA and do not specifically bind to any particular set of nucleotides (Figure 2B). We find that 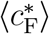 value is three orders of magnitude less compared to the 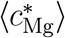 value implying the preferential exclusion of anion around the RNA (Figure 2A,B). Ion counting experiments^1,55^ and previous molecular dynamics simulation studies^2^ also support anion-exclusion from the ionic atmosphere surrounding the nucleic acids. The F^−^ binding to the FAD is primarily governed by the correct orientation of LBD-nucleotides where the three Mg^2+^ bind to constitute the cationic LBD (LBD-3Mg) that captures the F^−^ from solution (Figure 1A). At low [Mg^2+^] (= 1 mM), we found only 9 instances where a F^−^ is transiently bound to the aptamer, and the LBD is partially folded in all these conformations. In 8 out of 9 instances, the F^−^ is bound to the LBD with the assistance of only one Mg^2+^ (Figure S8A). For [Mg^2+^] (= 4 mM), the number of instances where one F^−^ bound to the LBD increased to 21. Out of 21, in 7 instances two Mg^2+^ capture a F^−^ to facilitate its binding to the partially folded LBD (Figure S8B). The number of F^−^ binding instances increased to 29 for [Mg^2+^] = 8mM. We found 15 instances where two Mg^2+^ capture a F^−^ and in 4 such instances the aptamer is in native-like folded conformations with the fully-folded LBD (Figure S8C). We found only one instance with three Mg^2+^ capture one F^−^ at LBD and the aptamer is in *holo*-form (Figure S8D). However, the average life-time of all the F^−^ bound conformations observed in the simulations using the coarse-grained RNA model is ⪅ 12.5 ps and it is not a stable state. The main conclusion from the simulations using the coarse-grained RNA model is that with the increase in [Mg^2+^], one F^−^ captured by two Mg^2+^ bound to the cationic LBD is the dominant binding mode for F^−^. The coarse-grained simulations also show that for the high [Mg^2+^] FAD can populate *holo*-like conformations even in absence of F^−^ binding, competent to activate transcription (Figure S8D).

The transient nature of F^−^ binding to the cationic LBD even at high [Mg^2+^] (≥ 5 mM), can be attributed to the lack of explicit solvent molecules in the coarse-grained simulations. We hypothesized that the lack of hydration shells around the Mg^2+^ prevented the binding of more than two Mg^2+^ in the fully formed LBD due to the enhanced electrostatic repulsion among the Mg^2+^. It is known that the lack of water mediated interactions can destabilize the binding of three Mg^2+^ and F^−^ in the LBD^3,31^ (Figure 1A). Furthermore, due to the lack of rigid hydration shell around the Mg^2+^, the cationic LBD is distorted and the Mg^2+^ cannot effectively counterbalance the electrostatic repulsion among the negatively charged phosphate beads of LBD-nucleotides. This resultant electrostatic repulsion among the negatively charged phosphate beads prevents the approach and binding of F^−^ to the LBD.

### Role of Mg^2+^ and Water in F^-^ Binding to the LBD

The coarse-grained simulations showed that the number of events where F^−^ binds to the LBD increased with [Mg^2+^]. At high [Mg^2+^], the dominant F^−^ binding mode to the FAD is one F^−^ captured by two Mg^2+^ bound to the LBD. We performed all-atom molecular dynamics simulations to probe the role of water in Mg^2+^ mediated F^−^ binding to the LBD. We performed six independent all-atom simulations starting from the FAD crystal structure (PDB: 4ENC)^3^ but without any Mg^2+^ and F^−^ bound to the LBD. This system is referred to as AA[0,0] (Table 1). The bulk concentration of both Mg^2+^ and F^−^ is 55 mM, and the concentration of K^+^ is 150 mM in the simulation box.

In two out of six AA[0,0] simulations we find that during the three step equilibration (see Methods), one Mg^2+^ water solvated ion which was closer to the phosphate oxygen atom of nucleotide G5 located in the vicinity of LBD formed a direct interaction (inner-sphere binding) with the phosphate oxygen atom of nucleotide G5. We treated the two systems where we have observed this as one Mg^2+^ bound near the LBD and referred it as AA[1,0] (Table 1). We have four independent simulations using system configuration AA[0,0]. The cumulative simulation time for both AA[0,0] and AA[1,0] configuration is tabulated in Table 1. The AA[0,0] and AA[1,0] system configurations enabled us to answer the following questions: (a) whether the cationic LBD pocket forms first due to the binding of three Mg^2+^ followed by F^−^ binding in the pocket? (b) how does F^−^ approach and bind to the LBD? (c) what is the configuration of the bound Mg^2+^ and F^−^ cluster in the LBD? and (d) what is the role of water in F^−^ binding to the LBD?

To track F^−^ binding to the LBD, we computed the coordination number for each of the F^−^ with the LBD RNA-atoms 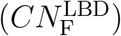, and the distance of each F^−^ from the center of mass of the LBD RNA-atoms 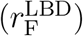 (see Methods) (Figure 3A and 4A). We consider an F^−^ is bound to the LBD if 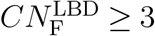 and 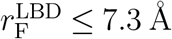. We identified the Mg^2+^ that facilitates F^−^ binding by computing the distance between all Mg^2+^ with the bound 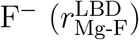 (Figure 3B and 4B). The Mg^2+^ that facilitated F^−^ binding are identified using the criteria 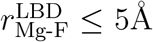.

**Figure 3:**
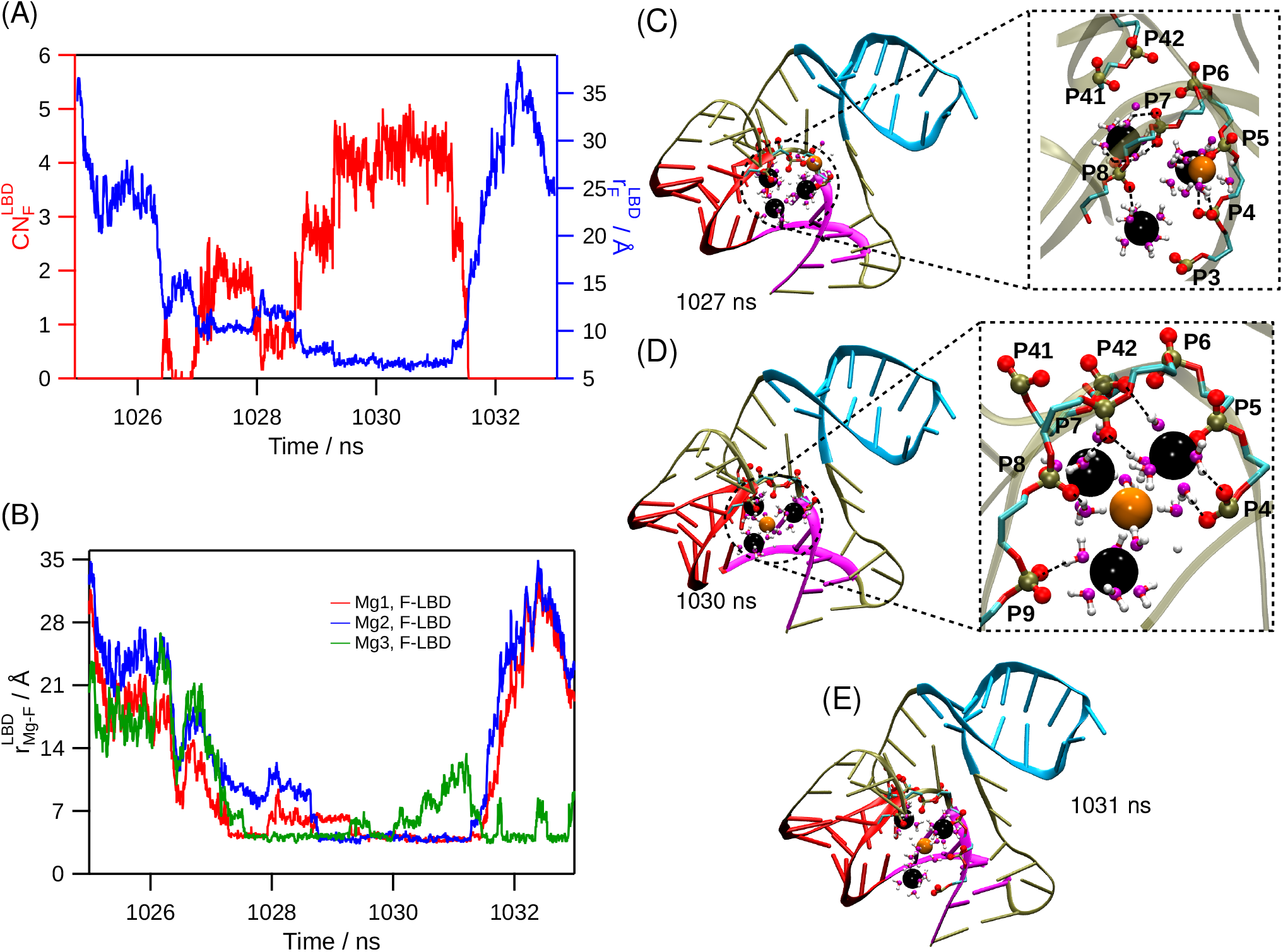
Kinetic pathway for the binding mode (I) obtained from the AA[1,0] simulations – where the three Mg^2+^ bind to the LBD-nucleotides to form the cationic pocket. The F^−^ later approaches the LBD and binds to the three Mg^2+^ ions. (A) Distance between the F^−^ ion, which binds to the LBD, and the center of mass of the LBD RNA-atoms is plotted in blue (right axis). The coordination number of the same F^−^ with the LBD RNA-atoms is plotted in red (left axis). (B) Distances between the bound F^−^ and the three Mg^2+^ are plotted. Distances for only those Mg^2+^ which are within water-mediated binding distance, 5 Å from the bound F^−^ are plotted. In panels (C) to (E), different stages of F^−^ binding to the LBD are shown. FAD is shown in cartoon representation. Pseudoknot PK, helices S_1_ and S_2_ are shown in magenta, red and cyan, respectively. Mg^2+^, F^−^, phosphorus and phosphate oxygen are shown as black, orange, tan and red spheres, respectively. Water oxygen and hydrogen are shown as magenta and white colored balls, using ball-and-stick representation, respectively. The backbone of LBD-nucleotides and nucleotide G5 are shown using stick representation. Cyan and red in the stick representation denote carbon and oxygen atoms, respectively. The nucleotide number (*N*_N_) of the phosphates and nucleobase are provided in the annotation. The snapshot simulation time is in the annotation of each panel. (C) F^−^ approaches the LBD with three Mg^2+^ bound to the LBD. (D) F^−^ interacts with the three Mg^2+^ bound to the LBD through water-mediated interaction to form the 3Mg-W-1F cluster. (E) F^−^ unbinding from the 3Mg-W-1F cluster.

In AA[0,0] simulations we found only one instance where F^−^ is captured by three Mg^2+^ to form water mediated 3Mg-W-1F cluster. In this F^−^ binding event, three Mg^2+^ bind to the LBD during the course of the simulation and F^−^ later approaches the LBD from bulk to form the 3Mg-W-1F cluster (Figure S10). Two out of three Mg^2+^ of 3Mg-W-1F cluster are bound to the cavity of LBD through water mediated interaction with the phosphate O atoms of LBD-nucleotides G8 and G42 (Figure S10C). The third Mg^2+^ in this cluster is bound at the periphery of the LBD through water mediated interaction with the phosphate O atoms of nucleotide G5 and phosphate O atoms of LBD-nucleotides (U7 and G8) rendering itself exposed to the bulk (Figure S10C). All three Mg^2+^ interact to the bound F^−^ through water-mediated interaction (Figure S10D). We discussed the implication of this solvent exposed Mg^2+^ on the ionic assembly formation in the following section.

From the all-atom simulations with system configuration AA[1,0] we found four instances where F^−^ is bound to the cationic LBD composed of three Mg^2+^. We observed the following two binding modes: (I) the three Mg^2+^ bind to the LBD and form the cationic LBD pocket (Figure 3C) and later F^−^ from the bulk binds to this cationic LBD pocket (Figure 3D,E), and (II) two Mg^2+^ bound to the phosphate groups of the LBD-nucleotides as observed in the coarse-grained simulations (Figure 4C). The third Mg^2+^ forms an ion-pair with a F^−^ in the bulk (Figure 4D) and the ion-pair binds to the LBD (Figure 4E,F). In both the binding modes, one Mg^2+^ is directly bound (inner-sphere) to the phosphate group of G5 located in the vicinity of LBD nucleotide throughout the simulation. Water molecules satisfy rest of the five coordinating sites of this Mg^2+^.

**Figure 4:**
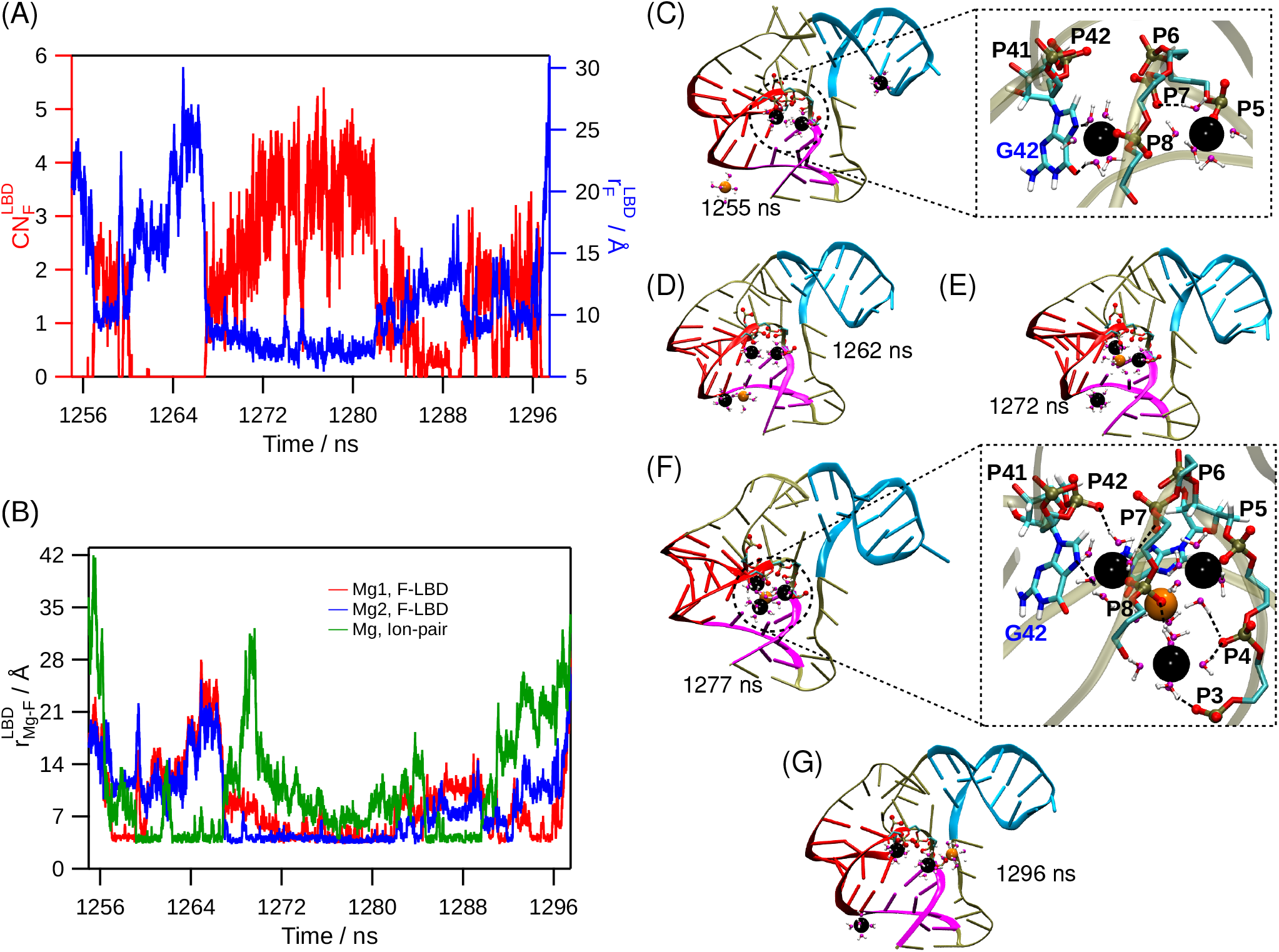
Kinetic pathway of F^−^ binding mode (II) obtained from the AA[1,0] simulations – where two Mg^2+^ bind to the LBD-nucleotides and third Mg^2+^ anchors the F^−^ to the LBD. (A) Distance between the F^−^ ion, which binds to the LBD, and the center of mass of the LBD RNA-atoms is plotted in blue (right axis). The coordination number of the same F^−^ with the LBD RNA-atoms is plotted in red (left axis). (B) Distances between the bound F^−^ and the three Mg^2+^ are plotted. Distance for only those Mg^2+^ which are within water mediated binding distance, 5 Å from the bound F^−^ are plotted. The distance between bound F^−^ and the Mg^2+^ of the ion-pair is plotted with green color. In panels (C) to (G) the different stages of F^−^ binding to the LBD are shown. FAD is shown in cartoon representation. Pseudoknot PK, helices S_1_ and S_2_ are shown in magenta, red, and cyan, respectively. Mg^2+^, F^−^, phosphorus and phosphate oxygen are shown in black, orange, tan and red colored spheres, respectively. Water oxygen and hydrogen are shown with magenta and white colored balls, using ball-and-stick representation respectively. The backbone of LBD-nucleotides, backbone of nucleotide G5 and nucleobase of G42 are shown with stick representation. Cyan, blue, red and white colors in the stick representation are used to denote carbon, nitrogen, oxygen and hydrogen atoms, respectively. The nucleotide number (*N*_N_) of the phosphates and nucleobase are provided in the annotation. The snapshot simulation time is in the annotation of each panel. (C)-(E) show the early stages of F^−^ binding to the cationic pocket and F^−^ approach to the LBD. (F) F^−^ bound conformation where 3 Mg^2+^ capture one F^−^ through water mediated interaction to form 3Mg-W-1F cluster. (G) F^−^ unbinding from the 3Mg-W-1F cluster.

In the binding mode (I) – the cationic pocket is preformed. Apart from the inner sphere bound Mg^2+^, the other two Mg^2+^ form water mediated interactions with the phosphate O atoms of nucleotides U7 and G8. The water solvated F^−^ from the bulk approaches the cationic pocket and binds to it. There is only one hydration layer between Mg^2+^ and the bound F^−^ in the 3Mg-W-1F cluster. Computational studies have shown that the energy barrier for Mg^2+^ to lose a single water molecule from its first solvation layer is high (≈ 18.8 kcal mol^−1^).^56^ The incoming F^−^ loses its water solvation shell and binds to the Mg^2+^ of cationic pocket interacting with the water hydrogen atoms of Mg^2+^ solvation shell to form 3Mg-W-1F cluster (Figure 3D).

In the binding mode (II) – the second Mg^2+^ ion approaches the LBD and binds to the N7 atom of nucleobase G42 through a water-mediated interaction (Figure 4C). The third Mg^2+^ forms a water mediated ion-pair with hydrated F^−^ in the bulk, and this ion-pair binds to the two prebound Mg^2+^ in the LBD forming a water-mediated cluster of 3 Mg^2+^ and 1F^−^ (3Mg-W-1F) (Figure 4D,F). The cluster 3Mg-W-1F is held together by the water-mediated interactions between the Mg^2+^ and F^−^ where water O atoms satisfy all the six coordinating sites of Mg^2+^, and the water H atoms interact with the F^−^ (Figure 4F). The 3Mg-W-1F cluster interacts with the LBD through water-mediated interactions between Mg^2+^ and backbone O atoms of the LBD nucleotides (phosphate groups of U7, G8, G42, and nucleobase of G42) and phosphate group of G5. The 3Mg-W-1F cluster has a average lifetime of ≈ 0.5 ns.

We observed that the third Mg^2+^, which forms a water-mediated ion-pair with F^−^ and forms part of the 3Mg-W-1F cluster, is not stable in the cluster (Figure 4E). This hydrated Mg^2+^ is weakly bound to the LBD compared to the other two Mg^2+^ because it cannot access the LBD cavity and is unable to form interactions with the LBD nucleotides (Figure 4F).

The incoming hydrated Mg^2+^ forms two water-mediated interactions with the phosphate O atoms of 5′ terminal nucleotides (G3 and C4) and LBD nucleotide (G8) (Figure 4F). These interactions are disrupted by the thermal fluctuations leading to the breaking away of the Mg^2+^ from the 3Mg-W-1F cluster (Figure 4G). The inability of the third Mg^2+^ to form a strong interaction with the phosphate backbone contributes to the two distinctive binding F^−^ modes to the LBD, as discussed earlier.

In all the F^−^ binding events obtained from both AA[0,0] and AA[1,0] simulations, the 3Mg-W-1F cluster is bound at the periphery of LBD (Figure 4C) and not at the core of LBD as in the crystal structure^3^ (Figure 1A). In the all-atom simulations, the water-mediated interactions between Mg^2+^ and the phosphate O atoms of LBD nucleotides result in weak binding of cationic pocket to the RNA. Even the interaction between Mg^2+^ and F^−^ is also water-mediated. In the crystal structure, Mg^2+^ and F^−^ interact directly and not through water-mediated interactions. In addition, all the three Mg^2+^ also have direct interaction with the phosphate O atoms (Figure 1A). To attain binding like the crystal bound state, each Mg^2+^ in the 3Mg-W-1F cluster should shed at least three water molecules from their first solvation shell. Due to the high energy barrier for Mg^2+^ losing the first water molecule from its first solvation shell (≈ 18.8 kcal mol^−1^),^56^ we did not observe this event in the all-atom unbiased simulations even on a time scale of ≥15 *μ*s. However, the magnitude of energy barrier strongly depends on the water and ion force field parameters.^57–59^

### Plausible Mechanism for Mg^2+^ and F^−^ Assembly in the LBD

All-atom simulations have shown that the water-mediated cluster 3Mg-W-1F is transiently bound to the RNA due to weak outer-shell interaction between the Mg^2+^ and phosphate groups of the LBD. For stable binding of the 3Mg-1F cluster at LBD, Mg^2+^ has to directly interact with the O atoms of the phosphate groups of LBD through inner-shell binding by shedding water molecules from its first solvation shell. A water molecule exchange from Mg^2+^ first solvation shell has a high energy barrier,^56,59–61^ which makes it highly unlikely that all the three Mg^2+^ in the 3Mg-W-1F cluster will simultaneously shed water molecules and assemble concertedly to form the 3Mg-1F cluster at LBD. The system can avoid this improbable event if the Mg^2+^ bind hierarchically and transition from outer to inner shell interaction in the LBD to construct the cationic pocket.

The longer Mg^2+^ is bound to the LBD with its water solvation shell, the higher is its probability of transitioning from an outer shell to an inner shell interaction. To show the plausibility of Mg^2+^ following the sequential assembly mechanism to form the 3Mg-1F cluster, we computed the average conditional lifetime and occupancy time of the three Mg^2+^ ions in the LBD. *τ*(*i, j*) is the average conditional lifetime that *i*^*th*^ Mg^2+^ ion with its water-mediated shell is bound to the LBD given that *j* number of Mg^2+^ are already prebound to the LBD. 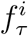 is the fraction of time at least *i* number of Mg^2+^ is bound to the LBD.

In the AA[0,0] system, the average lifetime of the first Mg^2+^ bound to the LBD is 27.31 ns as it is a water-mediated interaction. For it to bind stably to the LBD, it should undergo a transition from an outer to inner shell interaction, which we did not observe on our simulation timescale due to the high energy barrier (Table 2a). The lifetime of the second and third Mg^2+^ bound to LBD with their water solvation shells are even smaller, ≈ 1 ns, and 0.04 ns, respectively. Even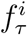 decreased with the increase in *i*, the number of Mg^2+^ bound to the LBD (Table 2b). Therefore it is unlikely that the three bound Mg^2+^ will transition from an outer to inner shell interaction with all of them bound to the LBD. A plausible mechanism is that the three Mg^2+^ transition from an outer to inner shell interaction in a hierarchical manner, and the following analysis supports it.

**Table 2:**
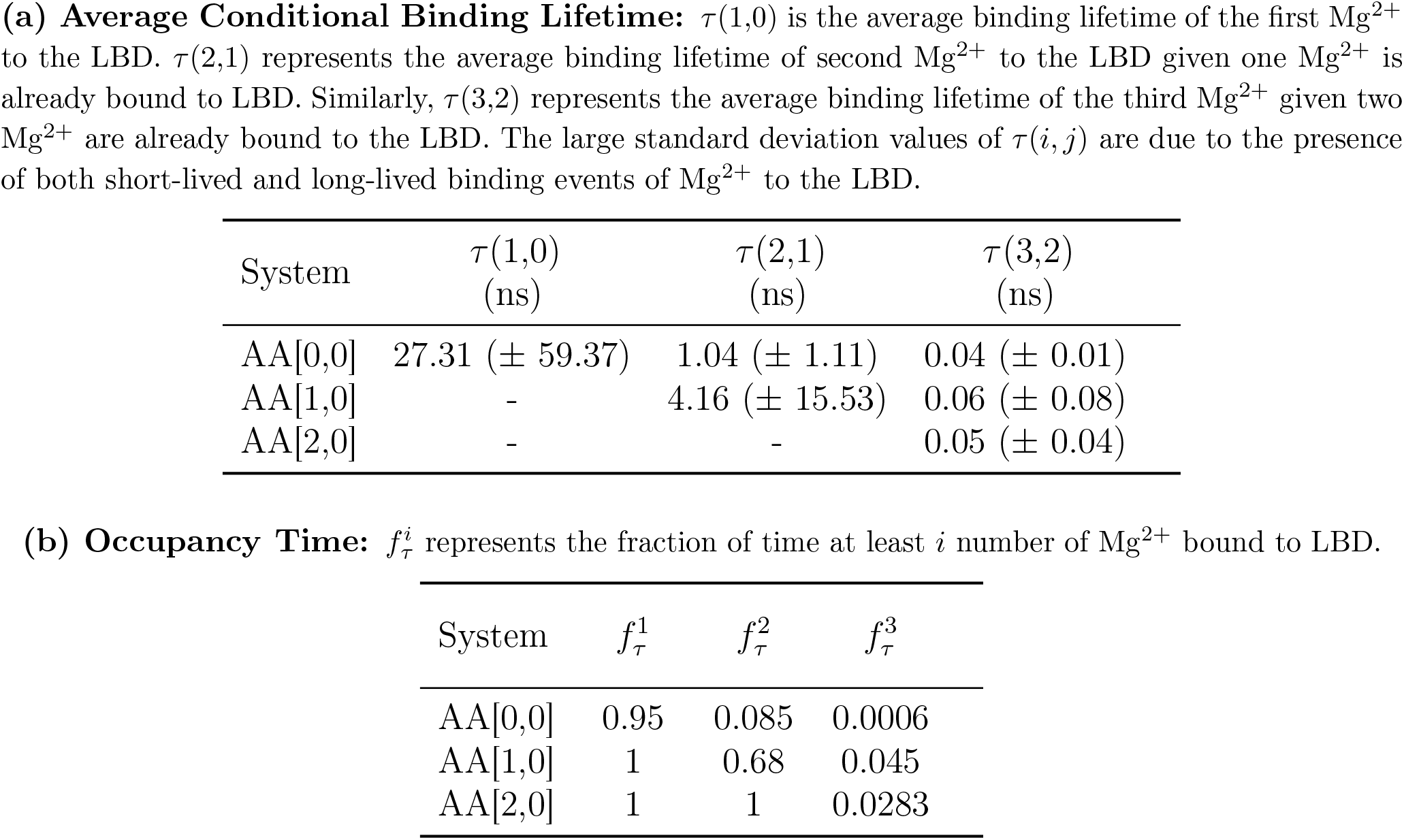
Mg^2+^ Binding Time with LBD in all-atom simulations.

| System | $\tau(1,0)$<br>(ns) | $\tau(2,1)$<br>(ns) | $\tau(3,2)$<br>(ns) |
| --- | --- | --- | --- |
| AA[0,0] | 27.31 (± 59.37) | 1.04 (± 1.11) | 0.04 (± 0.01) |
| AA[1,0] | - | 4.16 (± 15.53) | 0.06 (± 0.08) |
| AA[2,0] | - | - | 0.05 (± 0.04) |

| System | $f_{\tau}^1$ | $f_{\tau}^2$ | $f_{\tau}^3$ |
| --- | --- | --- | --- |
| AA[0,0] | 0.95 | 0.085 | 0.0006 |
| AA[1,0] | 1 | 0.68 | 0.045 |
| AA[2,0] | 1 | 1 | 0.0283 |

The values of *τ*(2, 1) and 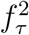 are larger for the AA[1,0] system compared to the AA[0,0] system (Table 2a and 2b). This indicates that when the first Mg^2+^ ion is bound to the LBD using inner shell interaction, then the frequency of the second Mg^2+^ ion binding to the LBD increases and also its bound lifetime, enhancing its probability to transition from an outer to inner shell interaction. This supports the mechanism that initially, the first bound Mg^2+^ transitions from outer to inner shell interaction in the LBD and facilitates the binding and transition of the second bound Mg^2+^ from outer to inner shell interaction later. However, the *τ*(3, 2) and 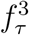 values are similar for all the systems indicating that even with the two Mg^2+^ already bound to the LBD with inner shell interaction (AA[2,0] system), binding of the third Mg^2+^ with its water solvation shell is probably hindered due to the electrostatic repulsion from the other two bound Mg^2+^ (Table 2a and 2b). As a result, we hypothesize that the efficient way for the third Mg^2+^ to approach the LBD with two prebound Mg^2+^ is, as an ion-pair with F^−^ to complete 3Mg-W-1F cluster, as observed earlier in the binding mode II. The third Mg^2+^ will later shed water from its solvation shell to transition from outer to inner shell interaction and complete the assembly of the 3Mg-1F cluster in the LBD.

### Fluoride Binding to the LBD Stabilizes Linchpin Hydrogen Bonds Critical for Transcription Initiation

We performed simulations of the FAD in AA[*ϕ*](*apo*) and AA[3,1](*holo*) configurations (Table 1) to identify the changes in the nucleotide fluctuations upon 3Mg-1F cluster binding to the LBD. The root mean square fluctuation (RMSF) computed at the nucleobase resolution shows that in the *apo* state, the nucleobases of LBD-nucleotides (A6, U7, G8, U41, and G42) are more flexible compared to the *holo* state (Figure S11B). The root mean square deviation (RMSD) of LBD-nucleotides shows that in the *holo* state, the three bound Mg^2+^ to the phosphate backbone of LBD-nucleotides stabilize their corresponding nucleobases in native-like orientation (Figure S11C).

We computed the dynamic cross-correlation, *C*_ij_ of FAD, for AA[*ϕ*](*apo*) and AA[3,1](*holo*) states to quantify the impact of 3Mg-1F cluster binding on the relative motion between the nucleobases that are critical for transmitting the ligand-binding signal to the expression platform. We plotted the difference in dynamical cross-correlation (Δ*C*_ij_) (see Eq. 9 in Methods) between the nucleobases *i* and *j* by subtracting *C*_ij_ of the *holo* state (AA[3,1]) from the *apo* state (AA[*ϕ*]) (Figure 5). Δ*C*_ij_ shows that the following nucleotides become anti-correlated due to the absence of 3Mg-1F cluster binding to the LBD in the *apo*-form: (i) A40 and C47-A49, (ii) G8 and A40, (iii) A6 and U38 (Figure 5).

**Figure 5:**
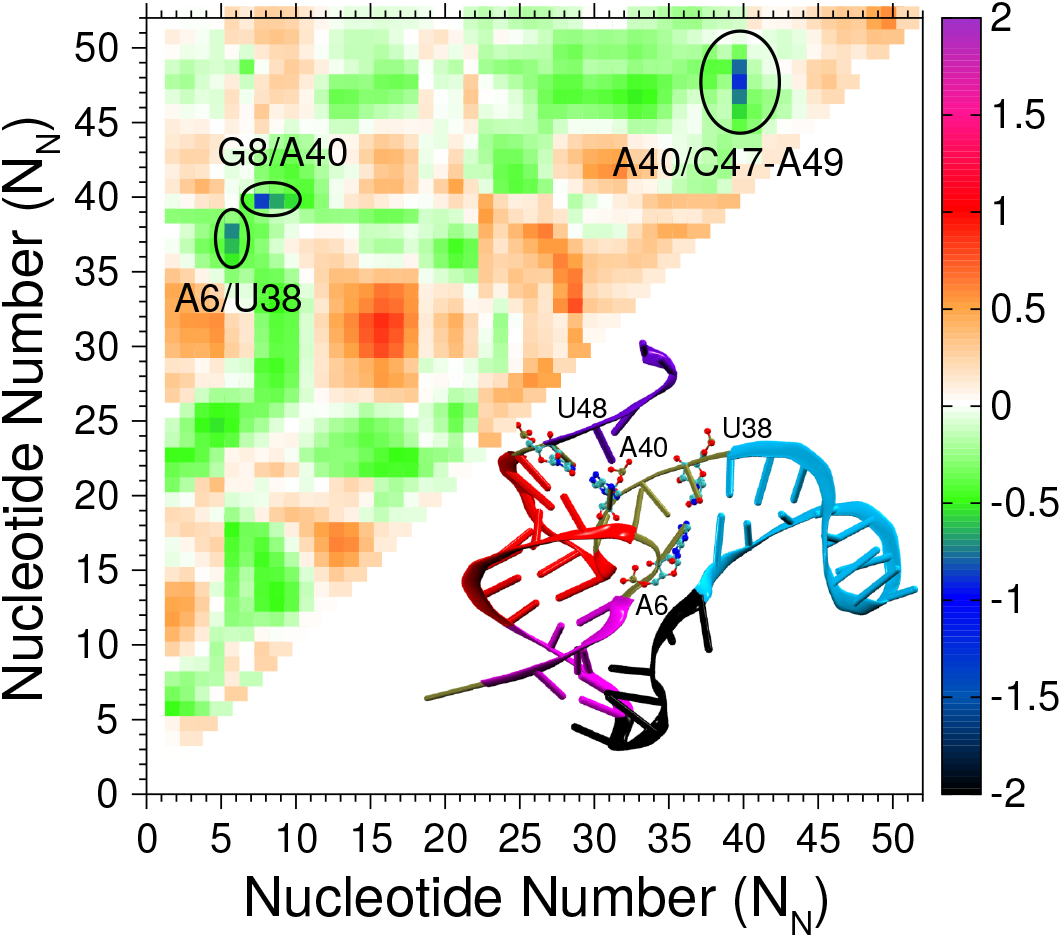
The heatmap of the difference in cross-correlations for nucleobases (Δ*C*_ij_) obtained by subtracting the *C*_ij_ of *holo* form (AA[3,1]) from the *apo* form (AA[*ϕ*]). Δ*C*_ij_ depicts the change in correlation between a pair of nucleobase beads upon binding of the 3Mg-1F cluster to the FAD. The FAD structure is shown in the inset using the cartoon representation. The linchpin hydrogen bond pairs are shown using the ball-stick model. Magenta, red, cyan, black, and violet colored regions represent nucleotides belonging to pseudoknot, helix S_1_, helix S_2_, PK-S_2_ and 3-TER, respectively.

We probed the structural changes in the RNA regions, which showed variations in the correlated motion (*N*_N_ = A6, G36-U38, A40 and U46-C50) upon 3Mg-1F cluster binding. The linchpin hydrogen bonds in the regions mentioned above break and only form in response to the 3Mg-1F cluster binding to the FAD (Figure S12B). The linchpin hydrogen bonds with reverse-Hoogsteen alignment (A40/U48) and reverse Watson-Crick alignment (A6/U38) are broken in the *apo* state (Figure S12B). This is in agreement with the experiments, which suggested that in the *apo* form, the linchpin nucleotide pair A40/U48 are not hydrogen-bonded to each other;^28^ instead, U48 forms hydrogen bonds with A64 located in the expression platform at the terminator helix.^3,29^ The terminator helix masks the polyU sequence from the RNA polymerase through strand invasion, and transcription is stopped.^3,29^ We find that the binding of the 3Mg-1F cluster to the FAD is imperative for forming both of the linchpin hydrogen bonds. Experiments also suggest that especially the formation of A40/U48 bond ensures the formation of the antiterminator helix, which exposes the polyU sequence to the RNA polymerase to activate the transcription (Figure S12B).^3,29^

## Conclusion

Enhanced levels of fluoride ion concentration in bacteria are detrimental to their survival. Fluoride riboswitch performs the critical function of maintaining fluoride ion homeostasis in bacteria. Its presence in multiple pathogenic bacteria makes it an ideal drug target for human pathogens. We studied the mechanism of fluoride ion recognition by the fluoride riboswitch. We showed that the riboswitch attains *holo*-like structure irrespective of F^−^ binding. However, it is not a globally stable structure in the absence of F^−^ binding. The native *holo*-form of the riboswitch becomes the most stable structure only upon F^−^ binding, which implies that it follows a conformational selection mechanism for the ligand binding. ^30^ The intermediates populated during riboswitch folding and cognate-ligand binding are potential targets for discovering new antibiotics. The computations support the sequential binding of the three Mg^2+^ ions to the LBD domain and transitioning from an outer to an inner shell interaction. The F^−^ ion most likely approaches the LBD by forming a water-mediated ion pair with the third Mg^2+^ ion.

For the riboswitch to control the transcription process, an interplay of the transcription speed, the folding time of the aptamer domain as the polymerase synthesizes it, and ligand binding to the aptamer domain are critical in determining the outcome of gene regulation.^28,29,62^ An exciting extension of this work would be to study the co-transcriptional folding of the fluoride riboswitch^28,29,63^ to explore at what stage of the aptamer domain synthesis does the fluoride ion binds and bifurcates the folding pathways, which either promote or inhibit the transcription terminator helix formation. However, this will be an extremely challenging computation. The three-dimensional structure of RNA can depend on the length of the RNA synthesized and transition into a different structure with the increase in length with time as the polymerase continues its synthesis. It will be challenging to perform these folding computations with the all-atom RNA models. To observe the folding of an RNA chain at a particular length, we have to reach micro to millisecond timescales. Further, we need to go beyond the second time scale to observe the transition into a different structure with the increase in RNA length. The development of coarse-grained RNA models which do not depend on the final crystal or NMR structure is essential for studying the transcriptional folding of RNA. A combination of coarse-grained and all-atom models can provide insights into this problem.

## Acknowledgement

GR acknowledges funding from the National Supercomputing Mission (MeitY/R&D/HPC/2(1)/2014). SK acknowledges research fellowship from Indian Institute of Science-Bangalore. We acknowledge National Supercomputing Mission (NSM) for providing computing resources of “PARAM Brahma” at IISER Pune, which is implemented by C-DAC and supported by the Ministry of Electronics and Information Technology (MeitY) and Department of Science and Technology (DST), Government of India.

