## Supplementary Information for "Mechanism of Mg^2+^ Mediated Encapsulation of an Anionic Cognate Ligand in a Bacterial Riboswitch"

### Supporting Information for "Mechanism of $Mg^{2+}$ Mediated Encapsulation of an Anionic Cognate Ligand in a Bacterial Riboswitch"

Sunil Kumar and Govardhan Reddy*

*Solid State and Structural Chemistry Unit, Indian Institute of Science, Bangalore, Karnataka, India 560012*

### Analyses

**Fraction of Native Contacts Among LBD-Nucleotides ($f_{\mathbf{LBD}}$)**: A pair of sites $i$ and $j$ belonging to the LBD-nucleotides (A6, U7, G8, U41 and G42) in the crystal structure are defined to have a native contact between them if $|i-j|>10$, and the distance between the sites $r_{ij}$ is less than 10 Å. The $f_{\mathrm{LBD}}$ for the $i^{th}$ conformation obtained from the simulations is computed using the equation

$$f_{LBD} = \frac{N^{i}_{\mathrm{LBD}}}{N^{\mathrm{cry}}_{\mathrm{LBD}}}, \tag{S1}$$

where $N^{i}_{\mathrm{LBD}}$ and $N^{\mathrm{cry}}_{\mathrm{LBD}}$ are the number of native contacts between the LBD-nucleotides present in the $i^{th}$ conformation and FAD crystal structure, respectively. The cut-off distance is the sum of largest distance between a pair of phosphate beads of LBD-nucleotides (7.8 Å) and the radius of phosphate bead (2.1 Å) to account for fluctuations.

**Fraction of Tertiary Contacts ($f_{\mathbf{TC}}$):** The nucleotides C18 - U23 (PK-$S_2$) form tertiary contacts with the nucleotides G1 - A6 of the pseudoknot (PK) labelled as $\mathrm{TC}_{\text{PK-S}_2}$. The total number of interactions in a given tertiary contact (TC) present in the FAD native state is equal to the number of native contacts between the sites belonging to the nucleotides involved in formation of that specific TC. The fraction of tertiary contacts $f_{\mathrm{TC}}$ for a particular TC is computed using the equation,

$$f_{TC} = \frac{N^{\mathrm{TC}}_{nc}(i)}{N^{\mathrm{TC}}_{\mathrm{nc}}(cry)}, \tag{S2}$$

where $N^{\mathrm{TC}}_{\mathrm{nc}}(i)$ and $N^{\mathrm{TC}}_{\mathrm{nc}}(cry)$ is the number of native contacts present between the sites belonging to the nucleotides involved in the formation of that specific TC in the $i^{th}$ conformation and the crystal structure, respectively. We label a TC as formed (F) if $\langle f_{\mathrm{TC}}\rangle \geq 0.5$ and as ruptured (R) otherwise.

**Fraction of Tertiary Stack Formation ($f_{\mathbf{TST}}$):** The average fraction of TST formation is computed using the equation

$$\langle f_{TST}\rangle = \left\langle \frac{N^{i}_{\mathrm{TST}}}{N^{\mathrm{cry}}_{\mathrm{TST}}} \right\rangle, \tag{S3}$$

where $N^{i}_{\mathrm{TST}}$ and $N^{cry}_{\mathrm{TST}}$ are the number of TST present in $i^{th}$ conformation and FAD crystal structure, respectively. $\langle\ \rangle$ denotes the average over all the conformations. A native tertiary stack (TST) is considered to be formed if its energy is lower than the thermal energy.

#### Local Folding Dynamics of the Unstructured Nucleotides

To identify the local regions in FAD, which exhibit dynamic transitions from the unfolded state to the folded state, we probed the changes in the following unstructured regions of the aptamer: (a) nucleotides between pseudoknot PK and helix $S_2$ (labeled as PK-$S_2$) and (b) nucleotides located at the $3'$ terminal (labeled as 3-TER nucleotides) (Figure 1A,B). We computed the fraction of local native contacts - (a) between PK-$S_2$ nucleotides and their neighboring nucleotides in the crystal structure (labelled as $N_{\text{PK-S}_2}$ nucleotides) ($f_{\text{PK-S}_2}$), (b) between 3-TER nucleotides and their neighboring nucleotides (labelled as $N_{\text{3-TER}}$) ($f_{\text{3-TER}}$).

The FES projected onto $f_{\text{PK-S}_2}$ shows that for $[\text{Mg}^{2+}] = 1$ mM, the aptamer conformations with the ruptured native contacts between PK-$S_2$ and $N_{\text{PK-S}_2}$, especially with the nucleotides $N_{\text{N}}$ = G1 - A6 are more stable (Figure S3B, S4). Upon increase in $[\text{Mg}^{2+}]$ ($\geq 4$ mM), the aptamer starts populating conformations with native contacts intact between PK-$S_2$ and $N_{\text{PK-S}_2}$ leading to the folded state ($f_{\text{NC}} \approx 0.7$) (Figure S3B). However, even in high $[\text{Mg}^{2+}]$, the unfolded state with the ruptured contacts between PK-$S_2$ and $N_{\text{PK-S}_2}$ is the most stable state.

The FES projected on $f_{\text{3-TER}}$ also shows similar trends with variation in $[\text{Mg}^{2+}]$ (Figure S3D). In low $[\text{Mg}^{2+}]$, the state corresponding to the aptamer conformations where the native contacts between 3-TER and $N_{\text{3-TER}}$ nucleotides are not formed is the stable state. With the increase in $[\text{Mg}^{2+}]$, aptamer conformations with intact native contacts around 3-TER are populated. However, even when $[\text{Mg}^{2+}] = 8$ mM, the conformations with the ruptured contacts are the most stable state. The stability of the aptamer folded state in the coarse-grained simulations increased with the increase in $[\text{Mg}^{2+}]$. Still, it is not the global minimum in the FES as the conformations with formed local native contacts at PK-$S_2$ and 3-TER nucleotides are not dominantly populated (Figure S5, S6 and S7). NMR experiments[1] on FAD also reported that the aptamer in *apo*-form populates unfolded conformations where the pseudoknot PK, helices $S_1$ and $S_2$ are fully-formed but the local contacts around regions PK-$S_2$ and 3-TER are ruptured.

#### FAD Structure is Stabilized by Tertiary Stacks

We tracked the formation of all six native tertiary stacks (TST), namely A6/G24, A6/G39, U7/G39, G8/A40, A40/A49, and G5/G42 present in the *holo*-form of the aptamer. In the folded state, the pseudoknot PK is in contact with the helix $S_1$ using the TST G5/G42. The rest of the TSTs are formed due to the intercalation of a nucleobase in between two other nucleobases. Nucleotide A40 intercalates between nucleotides G8 and A49 forming two TSTs - G8/A40 and A40/A49. Similarly, nucleotides A6 and G39 intercalate between nucleotides U7 and G24 forming three TSTs - U7/G39, G39/A6, and A6/G24 (Figure S1).[2]

For each TST, we computed the fraction of TST formation, $f_{\text{TST}}$ (see Eq. S3) to track their contribution to the folding thermodynamics of the FAD. The average fraction of TST formation $\langle f_{\text{TST}} \rangle$ shows that only G5/G42 remains formed for all [$Mg^{2+}$] irrespective of whether aptamer is in unfolded or folded state (Figure S12A).

The large fluctuations in $\langle f_{\text{TST}} \rangle$ show that these tertiary stacks are highly dynamic as they form and break with the aptamer folding and unfolding, respectively (Figure S12A). $\langle f_{\text{TST}} \rangle$ $< 0.5$ for the five TSTs (except G5/G42) signifies that the conformations with the ruptured TSTs are dominantly populated at all [$Mg^{2+}$]. In contrast, the large fluctuations at high [$Mg^{2+}$] ($>$ 4 mM) indicate that these TSTs form in the folded state. An NMR experiment[1] proposed that $F^-$ binding is indispensable for the stability of the *holo*-like conformation. We did not observe a stable $F^-$ binding event in the coarse-grained simulations. As a result, the aptamer prefers to stay unfolded with ruptured intercalated TSTs, irrespective of $Mg^{2+}$.

**Table S1: Parameters for the hydrogen bonding potential ($U_{HB}$).[3]**
**The potential is given by $U_{HB} = U_{HB}^0 \ \exp(-u)$,**
**where** $u = \Bigg[5(r-r_0)^2 + 1.5\Bigg\{(\theta_1-\theta_{1,0})^2 + (\theta_2-\theta_{2,0})^2\Bigg\} + 0.15\Bigg\{(\psi-\psi_0)^2 + (\psi_1-\psi_{1,0})^2 + (\psi_2-\psi_{2,0})^2\Bigg\}\Bigg]$

**(a) Canonical hydrogen bond parameters.**

Parameters are derived from coarse-grained structure of ideal A-form RNA helix.[?]

| base pair | $r_0$ (Å) | $\theta_{1,0}$ | $\theta_{2,0}$ | $\psi_0$ | $\psi_{1,0}$ | $\psi_{2,0}$ |
|---|---|---|---|---|---|---|
| A−U | 5.8815 | 2.7283 | 2.5117 | 1.2559 | 0.9545 | 1.1747 |
| C−G | 5.655 | 2.4837 | 2.823 | 1.3902 | 1.2174 | 0.7619 |

**(b) Nucleotide number of base-pairs forming canonical hydrogen bonds in FAD.**

| nucleotide($B_1$)–nucleotide($B_2$), (#H-bonds) | | | |
|---|---|---|---|
| 2G–17C,(3) | 3G–16C,(3) | 4C–15G,(3) | 5G–14C,(3) |
| 6A–38U,(2) | 8G–47C,(3) | 9A–46U,(2) | 10G–45C,(3) |
| 11G–44C,(3) | 12C–43G,(3) | 13C–42G,(3) | 24G–37C,(3) |
| 25C–36G,(3) | 26C–35G,(3) | 27C–34G,(3) | 28U–33A,(2) |
| 40A–48U,(2) | | | |

**Table S2: Nucleotide number of base-pairs forming base-sugar hydrogen bond in FAD.**

Parameters are derived from coarse-grained structure of FAD.

| B–S | $r_0$ | $\theta_{1,0}$ | $\theta_{2,0}$ | $\psi_0$ | $\psi_{1,0}$ | $\psi_{2,0}$ |
|---|---|---|---|---|---|---|
| C16–A21 | 8.3067 | 0.9611 | 2.0346 | 0.9149 | 1.4714 | 1.5074 |

**Table S3: Nucleotide number of base-pair forming $\pi - \pi$ tertiary stack in FAD.**

Parameters are derived from coarse-grained structure of FAD.

| $B_1$–$B_2$ | $r_0$ | $\theta_{1,0}$ | $\theta_{2,0}$ | $\psi_0$ | $\psi_{1,0}$ | $\psi_{2,0}$ |
|---|---|---|---|---|---|---|
| A40–A49 | 4.0759 | 1.7713 | 1.9281 | 3.1163 | 2.8246 | 2.7289 |
| U7–G39 | 4.0931 | 1.936 | 1.7906 | 2.4207 | -2.0544 | 0.2272 |
| A6–G39 | 4.742 | 1.8636 | 1.912 | -2.0749 | 0.4153 | 2.5458 |
| A6–G24 | 4.2645 | 2.2069 | 1.1365 | 1.1546 | 2.3472 | 3.034 |
| G8–A40 | 4.0578 | 1.5253 | 1.4924 | 1.9766 | -2.741 | -1.5106 |
| G5–G42 | 4.0988 | 1.9679 | 1.4632 | 0.5203 | 0.5705 | -2.8705 |

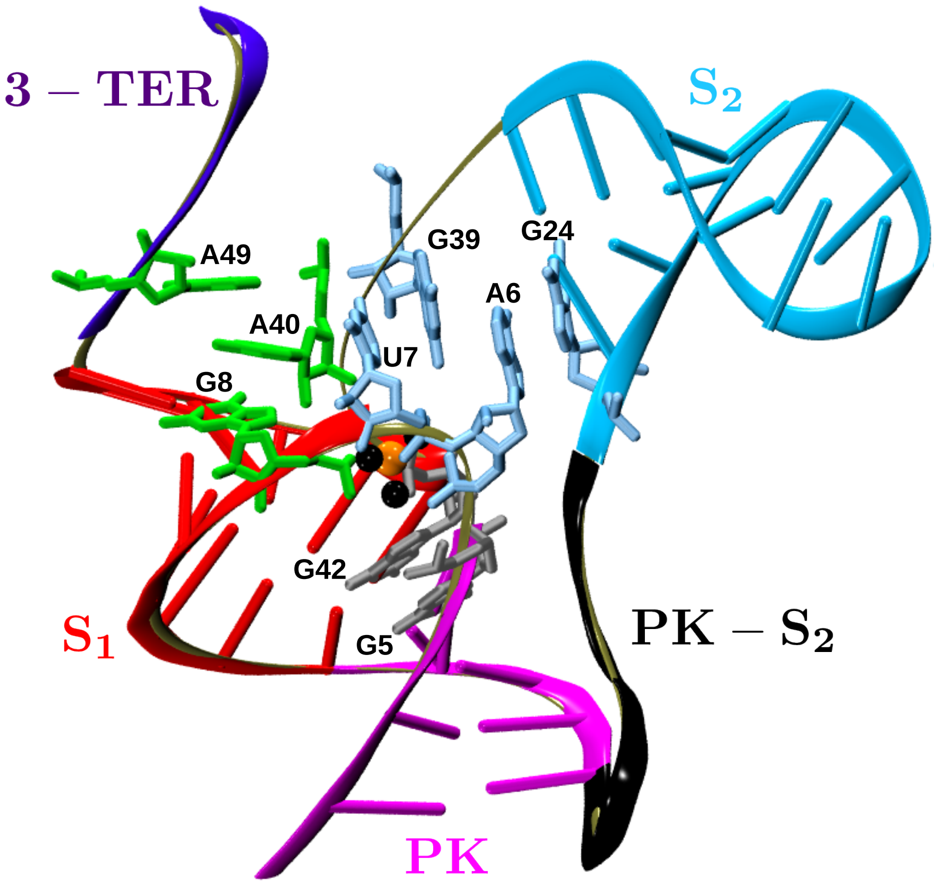


Figure S1: The six native tertiary stacks (TSTs) are shown on the three-dimensional structure of the FAD (PDB: 4ENC[2]). The RNA backbone is shown using a tan-colored cartoon representation. The pseudoknot PK, helices $S_1$ and $S_2$, unstructured nucleotides PK-$S_2$ and 3-TER are shown using magenta, red, cyan, black and purple colored cartoon representation, respectively. The two sets of intercalated tertiary stacks (TST) G8/A40; A40/A49, and A6/G24; A6/G39; U7/G39 are shown using green and steel blue colored bond representation, respectively. TST G5/G42 is shown using a gray-colored bond representation.

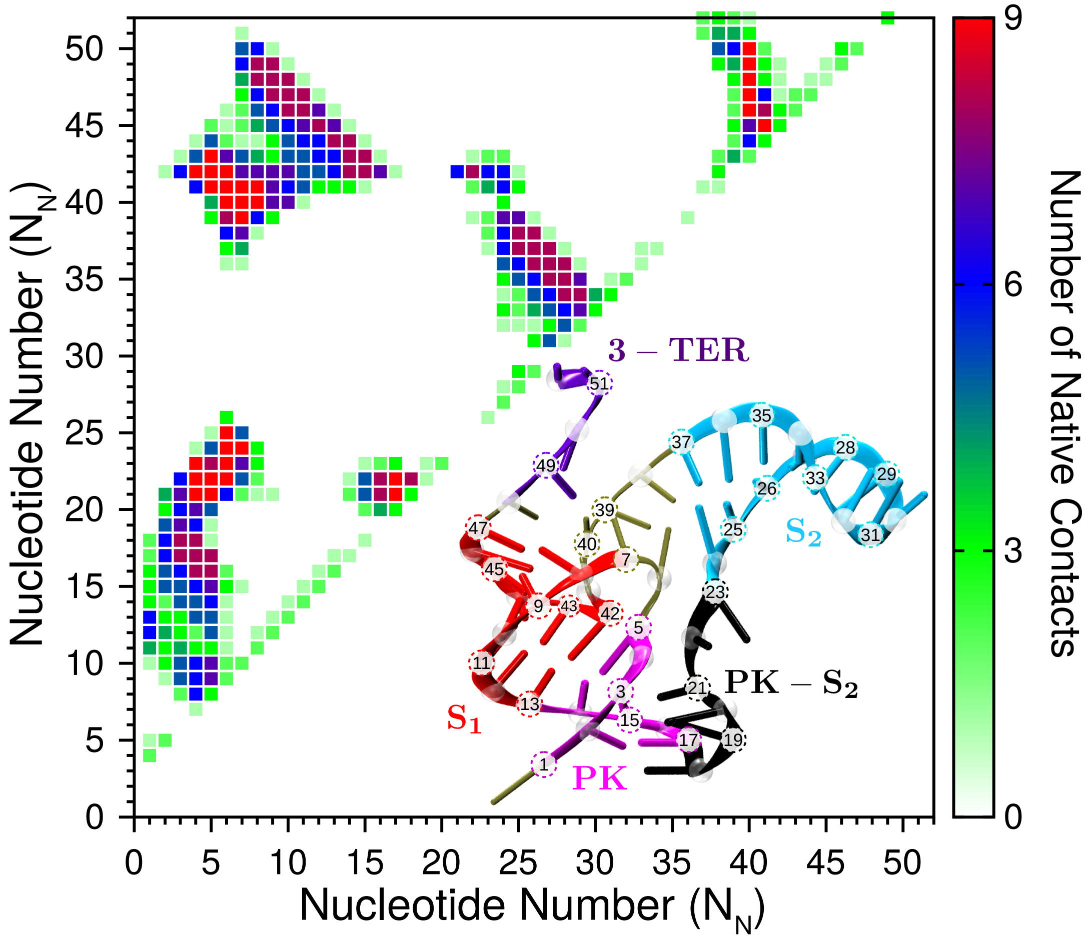


Figure S2: Native contact map of the FAD (PDB:4ENC)[2] at nucleotide resolution. The three-dimensional structure of the FAD is shown in the inset with marked nucleotide positions.

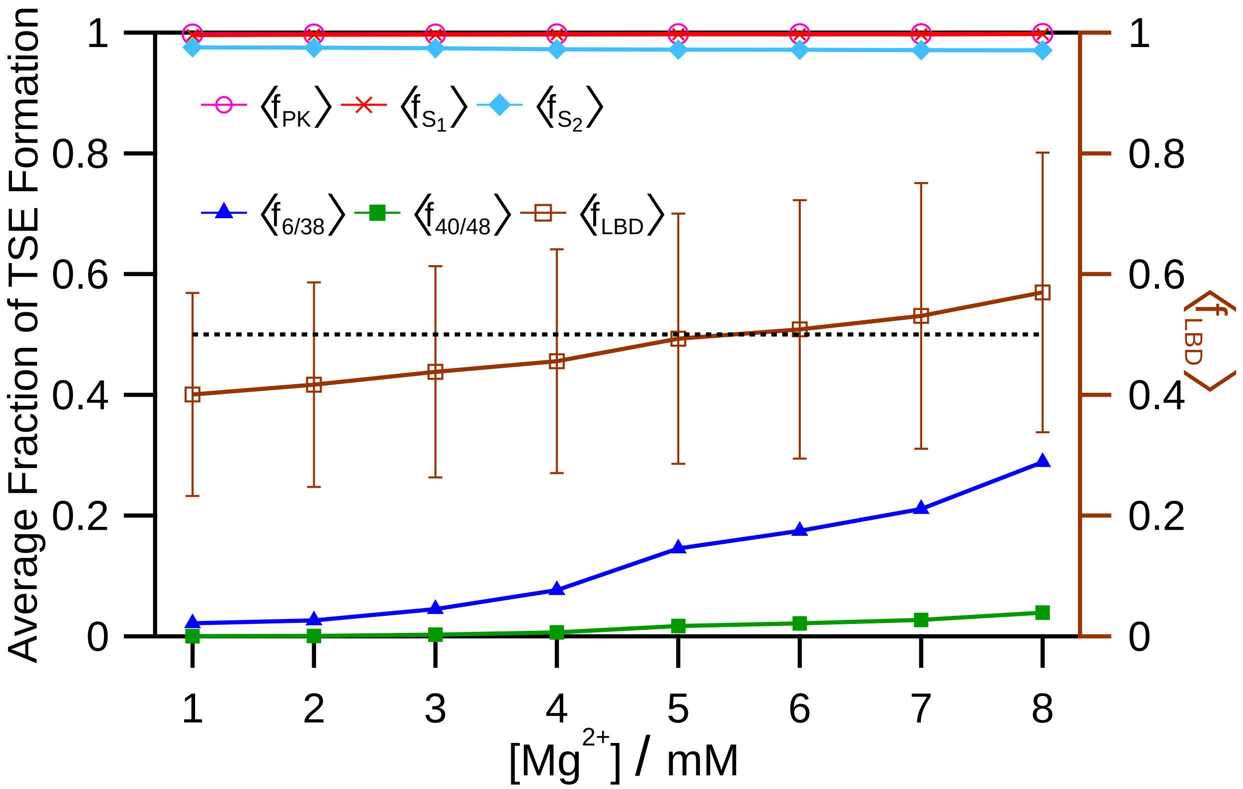


Figure S3: The average fraction of formation of tertiary structural elements (TSE), PK, $S_1$ and $S_2$ computed using the data from coarse-grained simulations (see Eq. 3 in Methods) are shown in magenta hollow circles, red crosses, and cyan rhombuses, respectively. The average fraction of formation for linchpin hydrogen bonds, A6/U38 ($\langle f_{6/38} \rangle$) and A40/U48 ($\langle f_{40/48} \rangle$) are shown in blue triangles and green squares, respectively. The average fraction of LBD formation (Eq. S1) is shown in brown hollow squares (right axis). The dotted black horizontal line is at 0.5 along both the left and right axes.

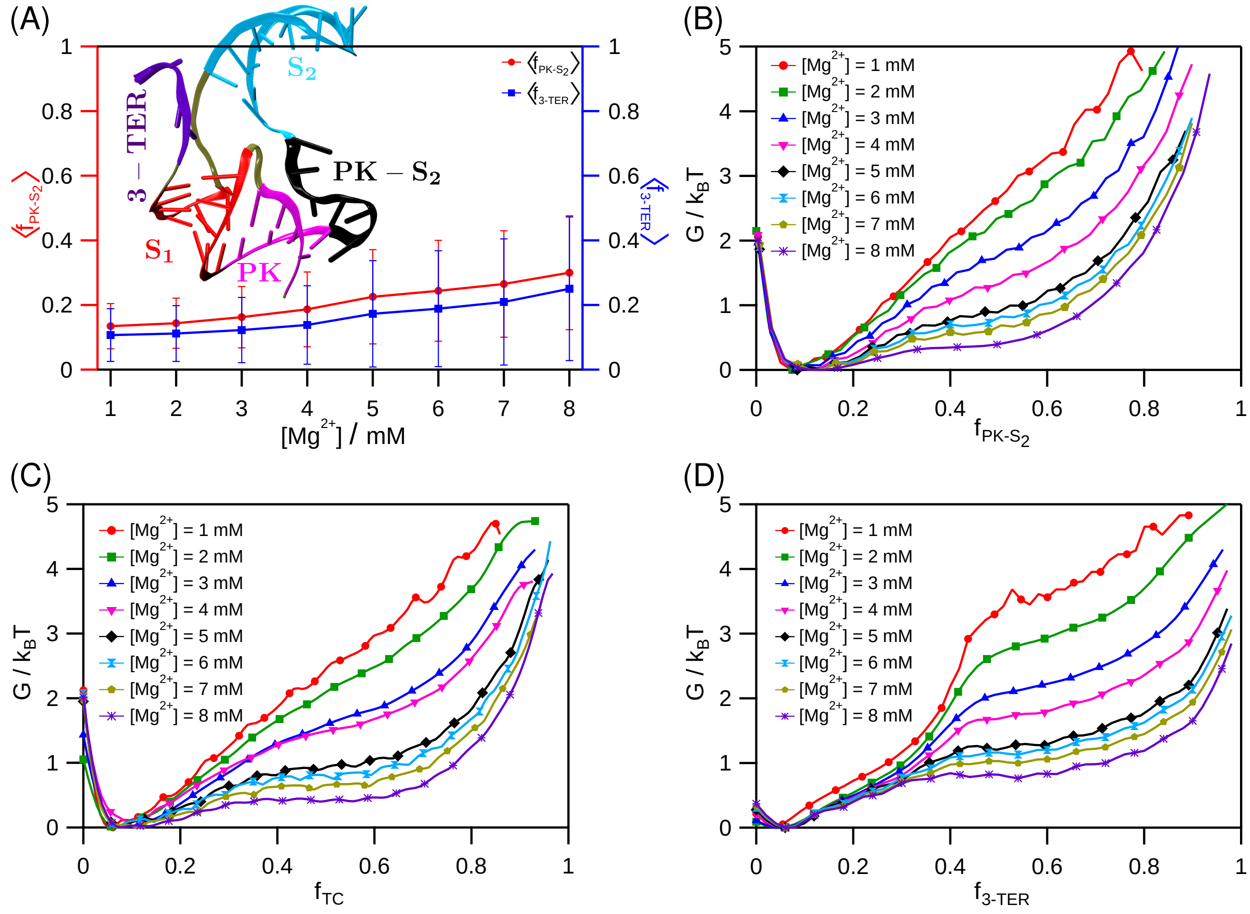


Figure S4: (A) The average fraction of native contacts between the PK-$S_2$ and $N_{\text{PK-}S_2}$ nucleotides ($\langle f_{\text{PK-}S_2} \rangle$) and average fraction of native contacts between the 3-TER and $N_{\text{3-TER}}$ nucleotides ($\langle f_{\text{3-TER}} \rangle$) computed using the coarse-grained simulation data. The FES projected on $f_{\text{PK-}S_2}$ (see Methods), fraction of tertiary contact (Eq. S2) and $f_{\text{3-TER}}$ (see Methods) for $[Mg^{2+}] = 1$ to 8 mM are plotted in panels (B), (C) and (D), respectively.

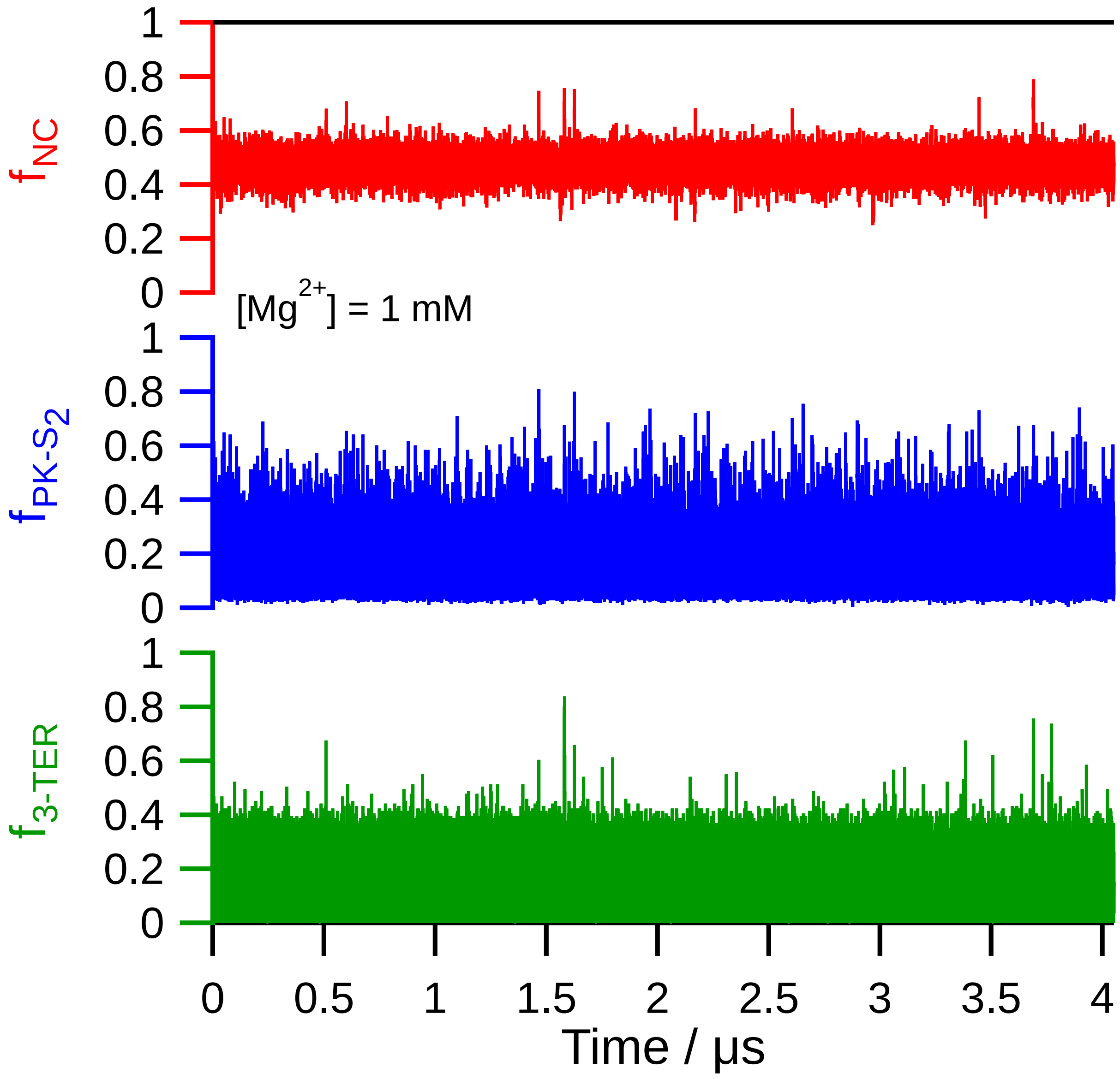


Figure S5: The fraction of native contacts ($f_{\mathrm{NC}}$), fraction of native contacts between PK-$S_2$ and $N_{\text{PK-S}_2}$ nucleotides ($f_{\text{PK-S}_2}$), and fraction of native contacts between 3-TER and $N_{\text{3-TER}}$ nucleotides ($f_{\text{3-TER}}$) are shown in top, middle and lower panels, respectively for $[\mathrm{Mg}^{2+}] = 1$ mM. We consider PK-$S_2$ to be formed if $f_{\text{PK-S}_2} > 0.5$. Similar criteria is used for $f_{\text{3-TER}}$ as well. Combined data for three independent trajectories is shown in the plots.

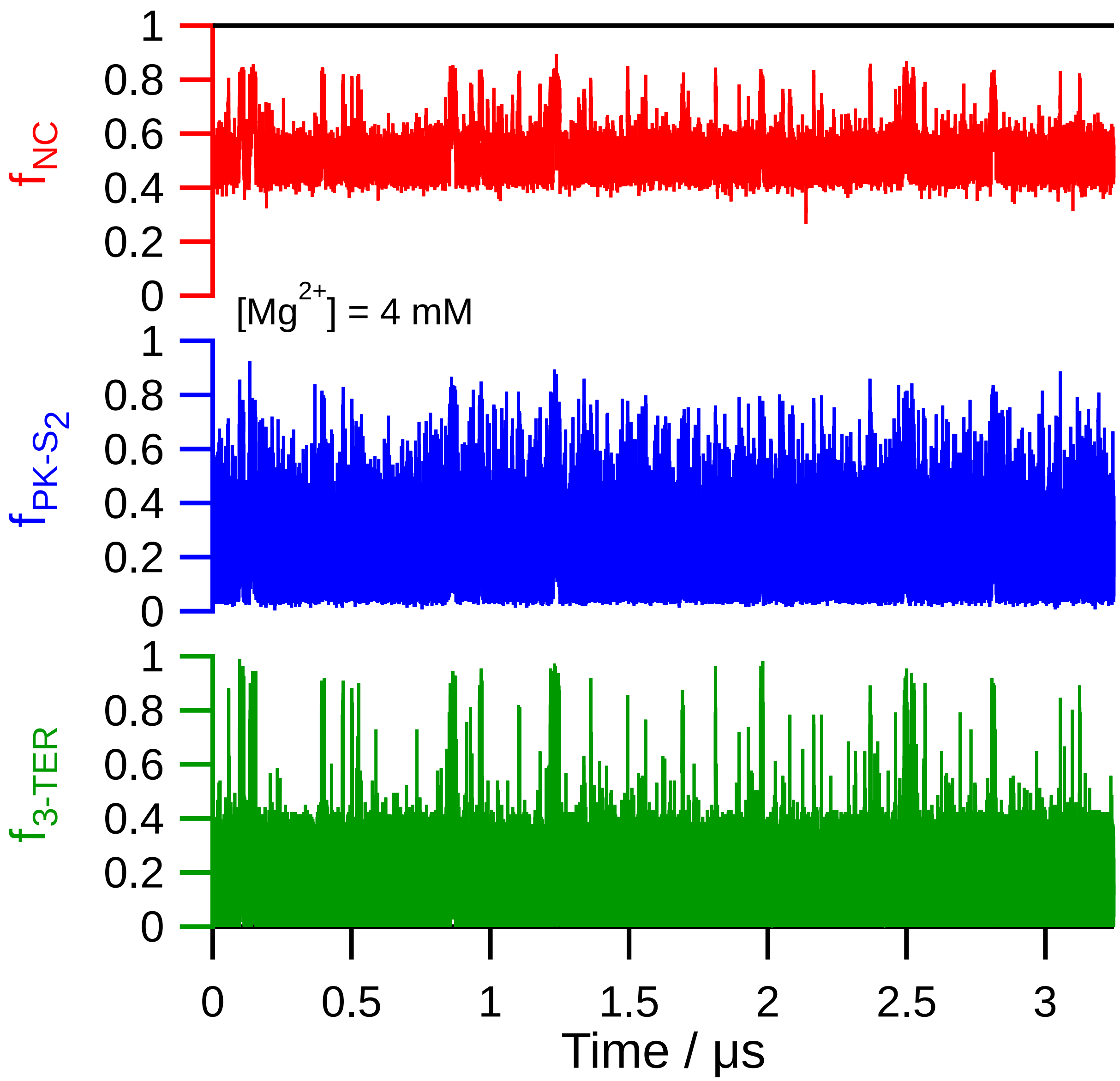


Figure S6: The fraction of native contacts ($f_{\mathrm{NC}}$), fraction of native contacts between PK-$S_2$ and $N_{\text{PK-}S_2}$ nucleotides ($f_{\text{PK-}S_2}$), and fraction of native contacts between 3-TER and $N_{\text{3-TER}}$ nucleotides ($f_{\text{3-TER}}$) are shown in top, middle and lower panels, respectively for $[Mg^{2+}] = 4$ mM.

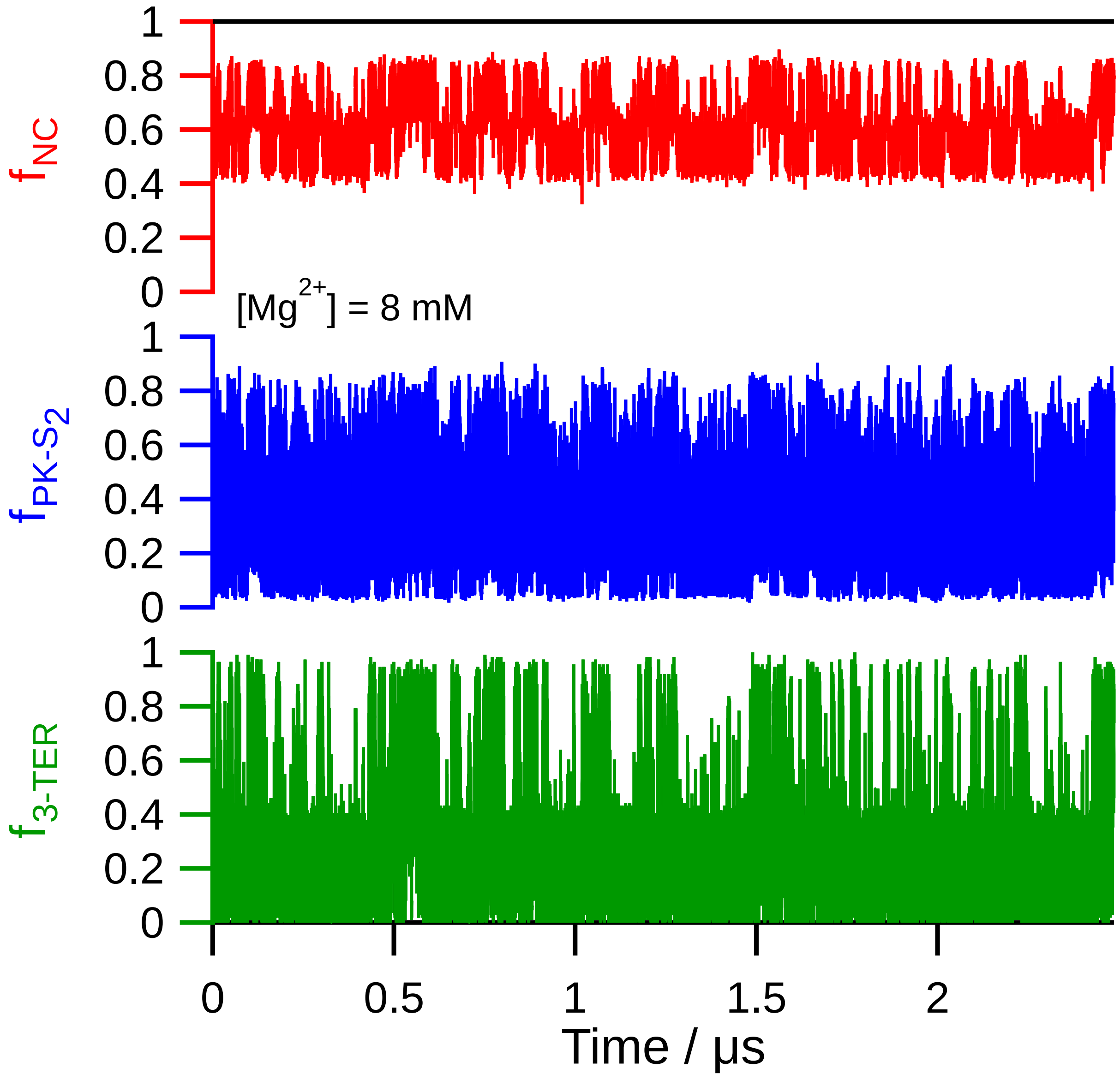


Figure S7: The fraction of native contacts ($f_{\mathrm{NC}}$), fraction of native contacts between PK-$S_2$ and $N_{\text{PK-S}_2}$ nucleotides ($f_{\text{PK-S}_2}$), and fraction of native contacts between 3-TER and $N_{\text{3-TER}}$ nucleotides ($f_{\text{3-TER}}$) are shown in top, middle and lower panels, respectively for $[Mg^{2+}] = 8$ mM.

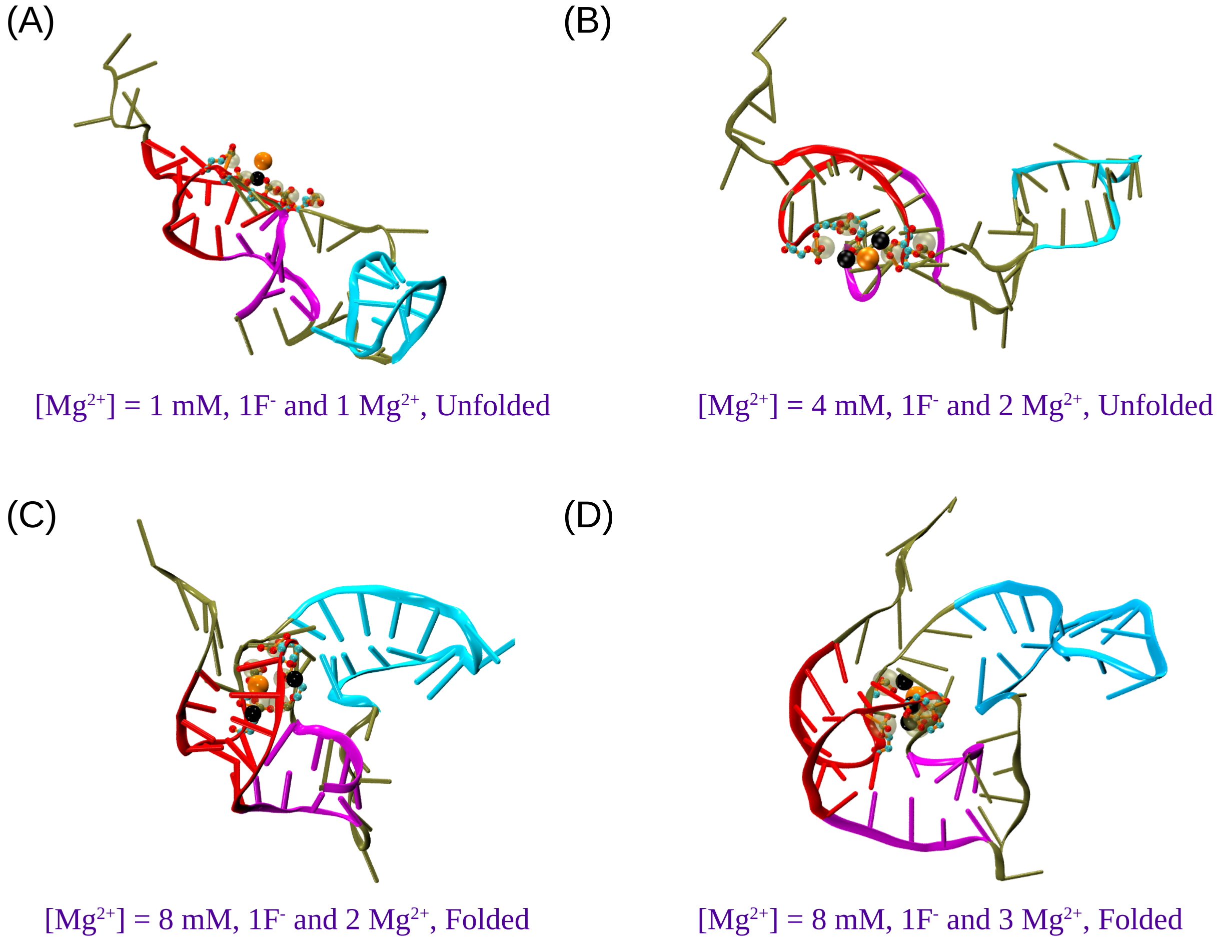


Figure S8: Representative snapshots from the coarse-grained simulations. The fluoride and magnesium ions bound to the LBD are shown as orange and black beads. (A) Unfolded FAD conformation at $[Mg^{2+}] = 1$ mM, (B) unfolded FAD conformation at $[Mg^{2+}] = 4$ mM, (C) native-like FAD conformation at $[Mg^{2+}] = 8$ mM, where two $Mg^{2+}$ and one $F^-$ are bound to LBD, (D) native-like FAD conformation at $[Mg^{2+}] = 8$ mM, where three $Mg^{2+}$ and one $F^-$ are bound to LBD.

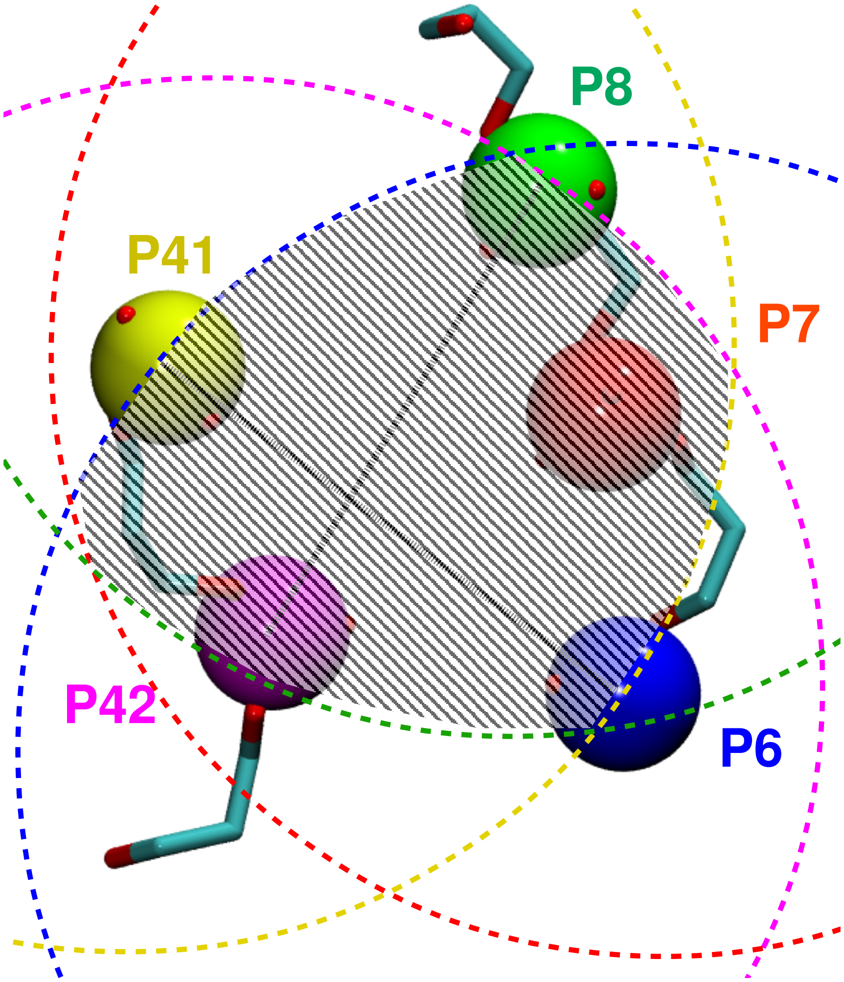


Figure S9: Schematic illustrating how an $Mg^{2+}$ bound to the LBD is identified in the all-atom simulations. The phosphorus atoms of the LBD nucleotides $N_{\mathrm{N}}$ = 6, 7, 8, 41, and 42 are shown as blue, red, green, yellow, and magenta colored beads, respectively. The two-dimensional representation of the sphere surfaces with radius $R_b$ are drawn as dashed lines with the phosphorus beads as the center with the same color as the beads. Any $Mg^{2+}$ within $R_b$ from all the phosphorus atoms of LBD-nucleotides (within the shaded volume) is considered to be bound.

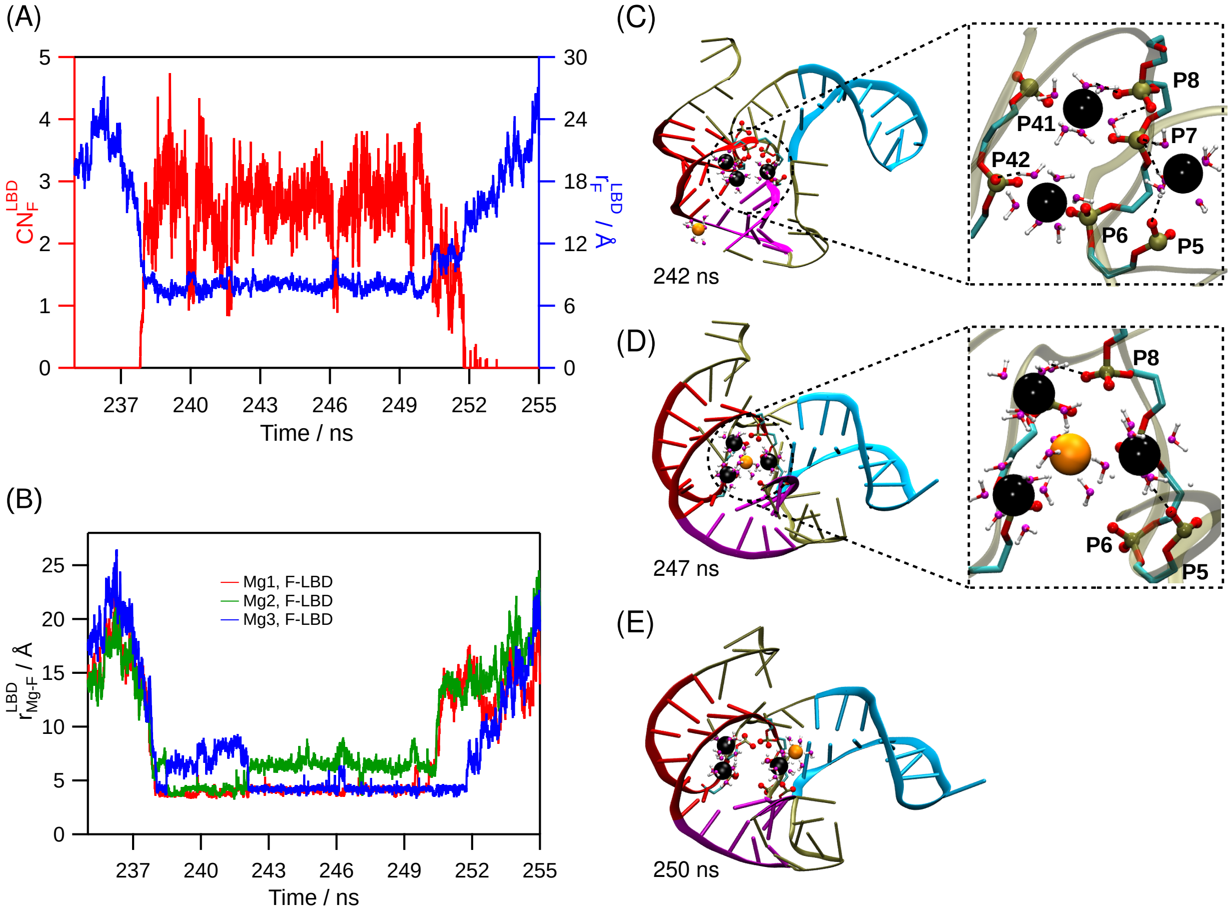


Figure S10: Kinetic pathway of $F^-$ binding obtained from the AA[0,0] simulations, where the three $Mg^{2+}$ bind to the LBD-nucleotides to form the cationic pocket. The $F^-$ later approaches the LBD and binds to the three $Mg^{2+}$ ions. (A) Distance between the $F^-$ ion, which binds to the LBD, and the center of mass of the LBD RNA-atoms is plotted in blue (right axis). The Coordination number of the same $F^-$ with the LBD RNA-atoms is plotted in red (left axis). (B) Distances between the bound $F^-$ and the three $Mg^{2+}$ are plotted. Distances for only those $Mg^{2+}$ which are within water-mediated binding distance, 5 Å from the bound $F^-$ are plotted. In panels (C) to (E), different stages of $F^-$ binding to the LBD are shown. FAD is shown in tan colored cartoon representation. Pseudoknot PK, helices $S_1$ and $S_2$ are shown in magenta, red, and cyan-colored cartoon representations, respectively. $Mg^{2+}$, $F^-$, phosphorus, and phosphate oxygen are shown in black, orange, tan, and red-colored spheres, respectively. Water oxygen and hydrogen are shown with magenta and white colored balls, using ball-and-stick representation, respectively. The backbone of LBD-nucleotides and nucleotide G5 are shown using stick representation. Cyan in the stick representation denotes carbon atoms. $N_N$ of the phosphates are provided in the annotation. The snapshot simulation time is in the annotation of each panel. Panels (C)-(E) show various stages of $F^-$ binding to the preformed cationic pocket. In panel-(C), $F^-$ approaches the LBD, with three $Mg^{2+}$ bound to the LBD. (D) $F^-$ interacts with the three $Mg^{2+}$ bound to the LBD through water-mediated interaction to form the 3Mg-W-1F cluster. (E) $F^-$ unbinding from the 3Mg-W-1F cluster.

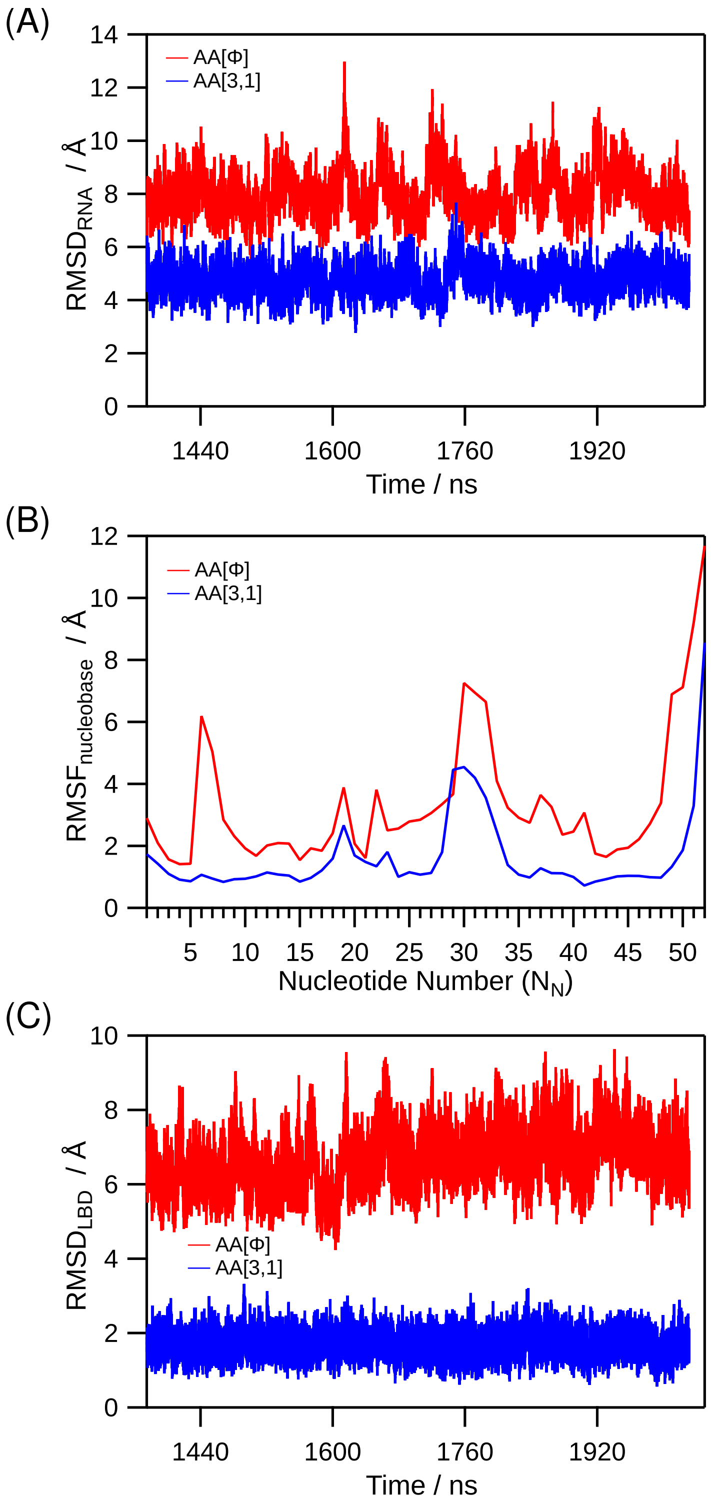


Figure S11: (A) RMSD of FAD for systems AA[$\phi$] (*apo*) and AA[3,1](*holo*) states are plotted with red and blue lines, respectively. RMSD is computed with respect to the energy minimized FAD crystal structure. (B) RMSF of the nucleobases for systems AA[$\phi$] (*apo*) and AA[3,1](*holo*) states are plotted with red and blue lines, respectively. (C) RMSD of the LBD for systems AA[$\phi$] (*apo*) and AA[3,1](*holo*) states are plotted with red and blue lines, respectively.

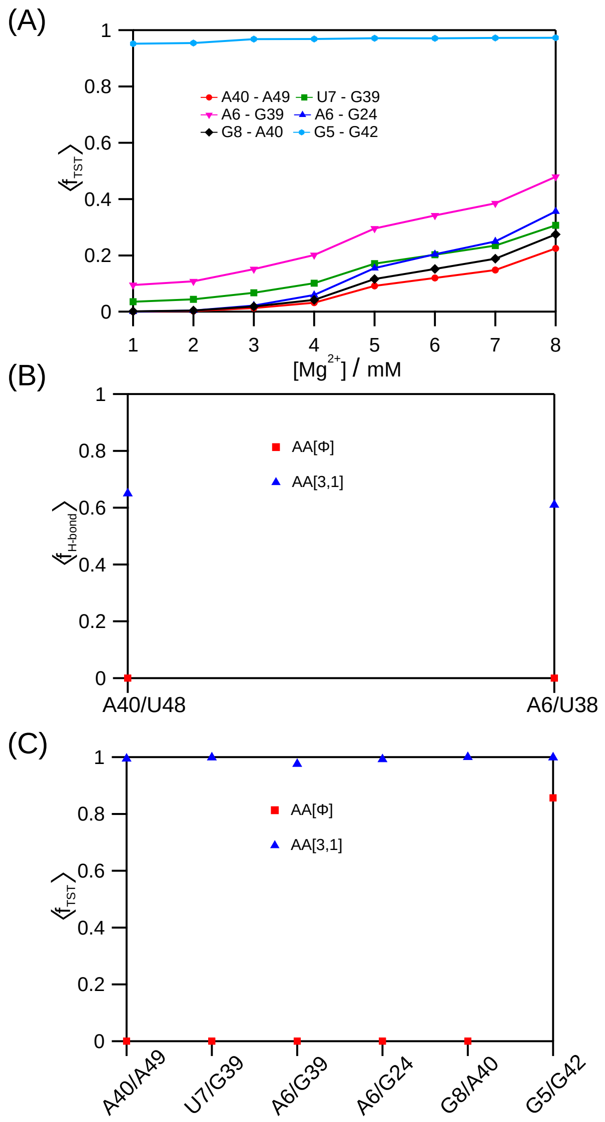


Figure S12: (A) The average fraction of formation of native tertiary stack, $\langle f_{\text{TST}} \rangle$ using the conformations obtained from CG simulations is plotted as a function of [$Mg^{2+}$]. $\langle f_{\text{TST}} \rangle$ for tertiary stacks between nucleotide pairs A40/A49, U7/G39, A6/A39, A6/G24, G8/A40 and G5/G42 are shown in red circle, green square, magenta inverted-triangle, blue triangle, black rhombus and cyan hexagon markers with lines, respectively. (B) The average fraction of native linchpin hydrogen bond formation, $\langle f_{\text{H-bond}} \rangle$ is plotted for the system setups AA[$\phi$] (*apo*) and AA[3,1] (*holo*) with red square and blue triangle solid markers, respectively. We used "hbonds" module of VMD[4] to compute the hydrogen bonds between the linchpins. We used "hbonds" module with 3 Å as distance cut-off between donor-acceptor atoms and with the default angle cut-off. (C) $\langle f_{\text{TST}} \rangle$ for all six TST is plotted for system setups AA[$\phi$] (*apo*) and AA[3,1] (*holo*) with red square and blue triangle solid markers, respectively. The name of the nucleotide pairs forming TST is given on the x-axis of the plot. Conformations obtained from the atomistic simulations were firstly converted to phosphate(P), Sugar(S) and nucleobase(B) using TIS-description. $\langle f_{\text{TST}} \rangle$ is then computed using Eq. S3.
